# Emx2 is an essential regulator of ciliated cell development across embryonic tissues

**DOI:** 10.1101/2024.01.04.574218

**Authors:** Thanh Khoa Nguyen, John-Michael Rodriguez, Hannah M. Wesselman, Rebecca A. Wingert

## Abstract

Cilia are hair-like organelles with vital physiological roles, and ciliogenesis defects underlie a range of severe congenital malformations and human diseases. Here, we report that the *empty spiracles homeobox gene 2 (emx2)* transcription factor is essential for cilia development across multiple embryonic tissues including the ear, neuromasts and Kupffer’s vesicle (KV), which establishes left/right axial pattern. *emx2* deficient embryos manifest altered fluid homeostasis and kidney defects including decreased multiciliated cells (MCCs), revealing that *emx2* is essential to properly establish several renal lineages as well. Further, *emx2* deficiency disrupted ciliogenesis on renal monociliated cells and MCCs, and led to aberrant basal body positioning in kidney cells. Interestingly, we discovered that *emx2* deficiency was associated with reduced expression of key factors which regulate prostaglandin biosynthesis: the transcriptional regulator *peroxisome proliferator-activated receptor gamma 1 alpha* (*ppargc1a*) and its downstream target *prostaglandin-endoperoxide synthase 1* (*ptgs1*), which encodes a crucial enzyme for production of the prostanoid ligand prostaglandin E2 (PGE_2_). Importantly, both ciliogenesis and renal fate changes were rescued when *emx2* deficient embryos were provided with PGE_2_ or transcripts encoding *ptgs1* or *ppargc1a*. Taken together, our findings reveal new essential roles of *emx2* in cilia development across several tissues, and identify *emx2* as a critical, novel regulator of prostaglandin biosynthesis during renal development and ciliogenesis.

## INTRODUCTION

Cilia are exquisite organelles that play crucial roles in ontogeny and physiology. Many vertebrate cells form a single cilium, while others form bundles of cilia and are termed multiciliated cells (MCCs). Proper ciliogenesis and ciliated cell fate choice are essential for diverse events ranging from body pattern formation to organ development, and defects in ciliated cell formation have catastrophic outcomes ranging from *situs inversus* to central nervous system disorders, retinopathies, liver and kidney diseases, among others. As such, continuing to elucidate the mechanisms that control ciliogenesis and ciliated cell type specification has broad reaching significance and medical relevance.

*Empty spiracles homeobox gene 2* (*emx2/Emx2/EMX2)* is a homeodomain transcription factor homologous to the empty spiracles *(ems)* gene in *Drosophila,* which is crucial for head development (Dalton et al., 1989; Walldorf and Gehring, 1992). In vertebrates, *emx2/Emx2/EMX2* is expressed during neuronal, auditory, olfactory and renal development (Simeone et al., 1992a,b; Gulisano et al., 1996; Miyamoto et al., 1997; Mallamaci et al., 1998; Tole et al., 2000; Cecchi, 2002; Galli et al., 2002; Kawahara and Dawid, 2002; Schubert and Lumsden, 2005; Brancaccio et al., 2010; Holley et al., 2010; Mariani et al., 2012; Jiang et al., 2017). Consistent with these sites of expression, previous studies uncovered a variety of phenotypes across tissues in animal models and humans deficient in *emx2/Emx2/EMX2.* For example, murine knockout of *Emx2* resulted in the disappearance of the kidney, ureters, gonads and genital tracts, and a lack of the Müllerian ducts (Pellegrini et al., 1996; Miyamoto et al., 1997). *Emx2* murine knockouts also display central nervous system defects such as in hippocampus development (Pellegrini et al., 1996; Yoshida et al., 1997; Tole et al., 2000). In humans, *EMX2* mutations have been associated with schizencephaly, a rare brain disorder (Brunelli et al., 1996; Faiella et al., 1997; Granata et al., 1997; Pang et al., 2008). Recently, single cell RNA-seq data from patients with autism spectrum disorders revealed disrupted expression of *EMX2* (Doostparast Torshizi and Wang, 2022), and patients with Mayer-Rokitansky-Küster-Hauser (MRKH), a rare syndrome involving malformation of Müllerian structures, have multiple variations in EMX2 sequence (Li et al., 2022). Most relevant to the present report, *emx2* deficiency has been linked to disrupted formation of several ciliated cell types. Specifically, *emx2/Emx2* is needed to establish planar cell polarity in mechanosensory hair cells located in the auditory and lateral line systems of zebrafish and mammals. In these hair cells, Emx2/EMX2 is requisite to properly position the basal body, from which the kinocilium will form during sensory cell differentiation, and alterations of this placement change the orientation of the stereociliary bundle (Holley et al., 2010; Jiang et al., 2017; Ji et al., 2018; Jacobo et al., 2019; Kozak et al., 2020). However, comparatively little is known about the role of *emx2* in the many other ciliated cell populations of the vertebrate embryo.

Here, we shed new light on the role of *emx2* in several ciliated organs. Using the zebrafish model, we detected the presence of *emx2* transcripts across developing ciliated cells and found that *emx2* deficiency disrupted cilia formation in the Kupffer’s Vesicle (KV), macula, neuromasts and the kidney. Further, *emx2* loss of function led to abnormal renal fluid flow and reductions in kidney MCC and distal late (DL) cell lineages—together, a suite of hallmarks that characterize developmental defects in prostaglandin signaling (Marra et al., 2019a; Chambers et al., 2020; Wesselman et al., 2023). Indeed, we found that *emx2* deficiency significantly reduced expression of key prostanoid biosynthesis regulators: the Ppargc1a transcription factor, encoded by *peroxisome proliferator-activated receptor gamma, coactivator 1 alpha (ppargc1a),* and the Ptgs1 cyclooxygenase enzyme, encoded by *prostaglandin-endoperoxide synthase 1* (*ptgs1*). Renal ciliated cell development was rescued in *emx2* deficient animals by provision of *ppargc1a* transcripts, *ptgs1* transcripts or exogenous treatment with the prostanoid prostaglandin E2 (PGE_2_). Interestingly, we also found that *emx2* deficiency led to aberrant basal body positioning in the kidney, echoing previous findings on the role of *emx2* in sensory cell polarity. Finally, PGE_2_ treatment rescued ciliogenesis in the KV of *emx2* deficient embryos and led to normal left/right axis formation, indicating that ciliary function was sufficiently restored to drive early patterning events—highlighting for the first time that prostaglandin signaling is vital to mediate ciliogenesis in non-renal cell types. In sum, our study has elucidated essential roles of *emx2* in ciliated cell development across multiple zebrafish embryonic tissues, and discovered that Emx2 influences prostaglandin biosynthesis to control renal and non-renal ciliated cell development.

## RESULTS

### *emx2* is essential for ciliated cell development across multiple zebrafish embryonic organs

Previous studies have linked *emx2/Emx2/EMX2* expression to developing ciliated tissues in several species. In the zebrafish, *emx2* has been reported to be expressed in the proximal region of the zebrafish embryonic kidney, known as the pronephros (Kawahara and Dawid, 2002). To further study where *emx2* is expressed in the zebrafish embryo, we performed whole mount *in situ* hybridization (WISH) in wild-type (WT) 28 somite stage (ss) embryos. Interestingly, we found *emx2* expression in several locations that contain ciliated cells, including the pronephros, otic vesicle, and nasal placode (Figure 1A). In the zebrafish kidney, there are two types of ciliated epithelia within the nephron functional units: monociliated cells and MCCs (Liu et al., 2007; Ma and Jiang, 2007; Nguyen et al., 2023). The populations of monociliated tubule cells in the zebrafish kidney are patterned into discrete functional segments known as the proximal convoluted tubule (PCT), proximal straight tubule (PST), distal early (DE) and distal late (DL) at the 28 ss (Supplemental Figure 1A) (Wingert et al., 2007). At this time, MCCs are located from the PCT to the DE in a “salt and pepper” fashion amongst the monociliated cells (Supplemental Figure 1A) (Marra and Wingert, 2016; Marra et al., 2019a,b). Within the pronephros, we noted that *emx2* transcripts were quite abundant within the region occupied by MCCs (Figure 1A). Therefore, we utilized double WISH using the MCC marker *odf3b* to assess whether MCCs or their neighboring epithelia expressed *emx2.* Interestingly, *emx2* transcripts colocalized with *odf3b^+^* cells by the 28 ss (Figure 1B). Furthermore, *emx2* transcripts were detected in the intercalated pronephric cells that were *odf3b^−^* at the 28 ss, thus indicating that the neighboring monociliated cells also expressed *emx2* (Figure 1B). We also detected the presence of *emx2* transcripts within the bilateral stripes of renal progenitors that give rise to monociliated and MCC progenitors from 5 ss to the 17 ss (Supplemental Figure 1C). Additionally, we observed *emx2* expression in the KV, a ciliated transient embryonic organ which initiates left/right (L/R) pattern formation of organs like the heart, thus has been likened to the mammalian node (Gokey et al., 2016; Dasgupta and Amack, 2016) (Supplemental Figure 1C). Taken together, the expression of *emx2* several ciliated organs suggested that *emx2* potentially plays a significant role in cilia development across the zebrafish.

**Figure 1.**
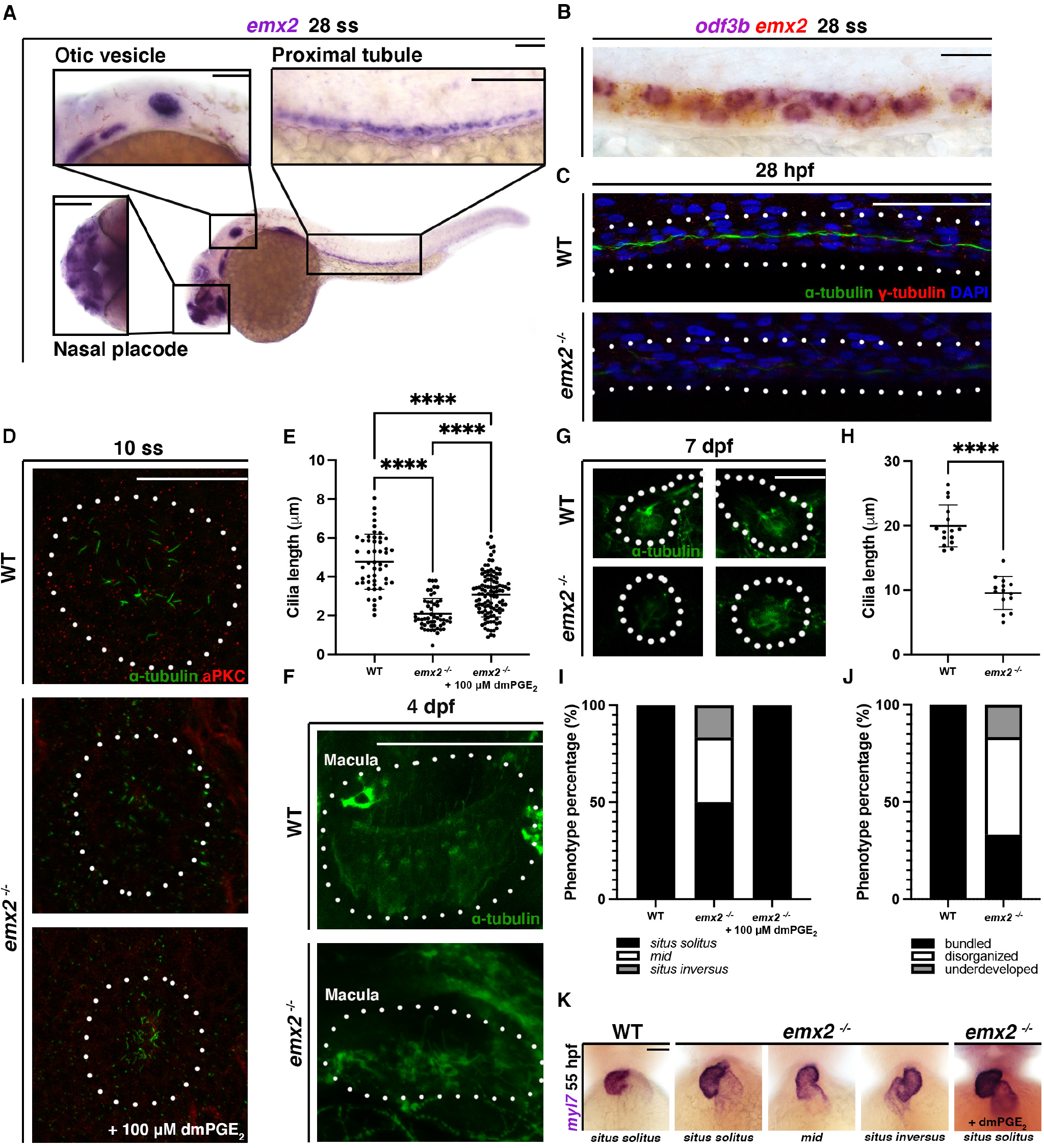
*emx2* is essential for cilia development in ciliated organs across the zebrafish embryo. (A) 28 ss WT embryo stained via WISH to illustrate *emx2* mRNA expression. Box is of approximate area of inset showing *emx2* expression in highly ciliated organs such as the otic vesicle, nasal placode and proximal tubule. Scale bar = 100 μm, otic vesicle and nasal placode inset = 50 μm, proximal tubule inset = 50 μm. (B) 28 ss WT embryo stained via WISH for *emx2* and an MCC marker (*odf3b)*. Scale bar = 50 μm. (C) 28 hpf whole-mount IF for acetylated α-tubulin (cilia, green), γ-tubulin (basal bodies, red), and DAPI (nucleus, blue) in the proximal pronephros of WT and *emx2^−/−^* embryos. Scale bar = 50 μm. (D) 10 ss whole-mount IF for acetylated α-tubulin (cilia, green) and anti-PKC (membrane boundary, red) in the KV of WT, *emx2^−/−^* embryos and *emx2^−/−^*embryos treated with dmPGE_2_. Scale bar = 50 μm. (E) KV cilia length between 10 ss WT, *emx2^−/−^* embryos and *emx2^−/−^* embryos treated with dmPGE_2_. (F) 4 dpf whole-mount IF for acetylated α-tubulin (cilia, green) in the macula of WT and *emx2^−/−^*embryos. Scale bar = 50 μm. (G) 7 dpf whole-mount IF for acetylated α-tubulin (cilia, green) in the neuromasts of WT and *emx2^−/−^*embryos. Scale bar = 25 μm. (H) Neuromast cilia length between 7 dpf WT *emx2^−/−^*embryos. (I) Phenotype percentage of neuromast phenotypes between WT and *emx2^−/−^*embryos. (J) Phenotype percentage of heart looping phenotypes between WT, *emx2^−/−^* embryos and *emx2^−/−^* embryos treated with dmPGE_2_. (K) 55 hpf WT, *emx2^−/−^* embryos and *emx2^−/−^*embryos treated with dmPGE_2_ stained via WISH using heart marker *myl7.* Scale bar = 50 μm. Data presented on graphs are represented as mean ± SD; * p<0.05, ** p< 0.01 ***p < 0.001 and ****p < 0.0001 (t-test or ANOVA).

To investigate the hypothesis that *emx2* is involved broadly in ciliogenesis, we examined cilia formation across tissues using the *emx2^el586/el586^* mutant line, referred to subsequently here as *emx2^−/−^*, which encodes a 10 basepair deletion that induces a frameshift after amino acid 74 (of 247) (Supplemental Figure 1B,E,F) (Askary et al., 2017). We performed whole-mount immunofluorescence (IF) to detect acetylated α-tubulin, which marks cilia (Wesselman et al., 2023). Compared to WT embryo controls, *emx2^−/−^* mutants displayed dramatic reductions in cilia formation within the pronephros segments occupied by MCCs (Figure 1C), as well as in distal segment regions where only monociliated cells are formed (Supplemental Figure 1D). Next, we utilized whole-mount IF to examine the KV organ during somitogenesis. At the 10 ss, *emx2*^−/−^ mutants exhibited reduced cilia length in the KV compared to WT embryos (Figure 1D,E). *emx2*^−/−^ mutants also displayed clear defects in otic vesicle development versus WT embryos, including a reduced and disorganized macula with shorter and disorganized cilia development (Figure 1F). Furthermore, using whole-mount IF at 7 days post fertilization (dpf), we investigated ciliogenesis in the neuromasts, which are small epithelial receptor organs sensory hair cells. Each sensory hair cell has a ciliary bundle composed of a single long kinocilium and many shorter stereocilia that sit to one side of the kinocilium. While WT embryos had normal neuromast structures with cilia bundled together, *emx2*^−/−^ mutants had disorganized cilia structures with unbundled cilia that appeared diminished in size (Figure 1G). In addition, we observed low α-tubulin intensity within neuromasts from some *emx2*^−/−^ embryos, suggesting the underdevelopment of neuromast cell cilia in these embryos. Quantification of cilia length and organization in WT and *emx2*^−/−^ revealed a statistically significant reduction in neuromast cell cilia length in the mutant embryos (Figure 1H) as well as large proportion of *emx2*^−/−^ animals with disorganized or underdeveloped cilia (Figure 1I).

We next explored whether changes in ciliary function might be consequent to one of these overt morphological defects in cilia development. To do this, we investigated whether changes in L/R patterning, which is driven by the KV motile cilia, were associated with *emx2* deficiency by examining cardiac looping. We conducted WISH with WT and *emx2*^−/−^ embryos at 55 hours post fertilization (hpf) using the cardiac marker *myl7* (Yelon et al., 1999). While all WT embryos displayed normal heart looping, some *emx2*^−/−^ embryos experienced randomization of heart looping, displaying *situs inversus* or *mid* phenotypes (Figure 1J,K). Taken together, these studies indicate that *emx2* plays several vital roles in ciliated cell development across the zebrafish embryo which have not been previously appreciated.

### *emx2* deficiency causes phenotypes associated with renal cilia defects

Given the required roles of *emx2* in ciliated sensory cells (Holley et al., 2010; Jiang et al., 2017; Ji et al., 2018; Jacobo et al., 2019; Kozak et al., 2020), and its co-expression with ciliated cells in the zebrafish pronephros, we next sought to further explore how loss of *emx2* function affects renal cilia development using several independent methods. In addition to the *emx2^−/−^* line, we developed an antisense knockdown model and an independent genetic model using CRISPR/Cas9. For the antisense strategy, we designed *emx2* splice-blocking morpholino oligonucleotides (MOs) to target the splice donor and splice acceptor junctions between exon 1 and exon 2 (Supplemental Figure 2A). Using RT-PCR and Sanger sequencing, we determined that the MOs resulted in the inclusion of intron 1, causing an in-frame stop codon, resulting in a truncated Emx2 protein (Supplemental Figure 2B,C). For an independent mutant model, we generated F0 mosaic *emx2* crispants by multiplexing multiple sgRNAs targeting three distinct regions of exon 1 (Supplemental Figure 3A). Crispant genotypes were confirmed using a T7 Endonuclease assay and Sanger sequencing (Supplemental Figure 3B,C), and we found that the *emx2* sgRNA multiplex was >90% penetrant in inducing mosaicism (Supplemental Figure 3D).

This suite of *emx2* loss of function models were then used to further explore the renal phenotypes associated with *emx2* deficiency, beginning with live morphology studies. At 48 hpf, the *emx2* morphant, *emx2* crispant and *emx2*^−/−^ embryos all exhibited pericardial edema and hydrocephalus, indicative of impaired kidney function (Figure 2A). We hypothesized that these *emx2* deficient phenotypes could be due to the lack of fluid flow resulting from changes in pronephros development, such as alterations in MCC development which frequently lead to the reduction or abrogation of luminal flow (Marra et al., 2019a; Chambers et al., 2020; Wesselman et al., 2023). To investigate this, we performed WISH analysis using the specific MCC marker *odf3b* in WT and *emx2* deficient embryos. *emx2* morphant, *emx2* crispant and *emx2*^−/−^ embryos had decreased *odf3b^+^*cells per embryo compared to WT controls (Figure 2B,C). Additionally, WISH analysis using the specific MCC marker *cetn4*, which, similar to *odf3b,* marks differentiating MCCs, revealed a statistically significant reduction in MCC number in *emx2* deficient embryos (Supplemental Figure 4A,D).

**Figure 2.**
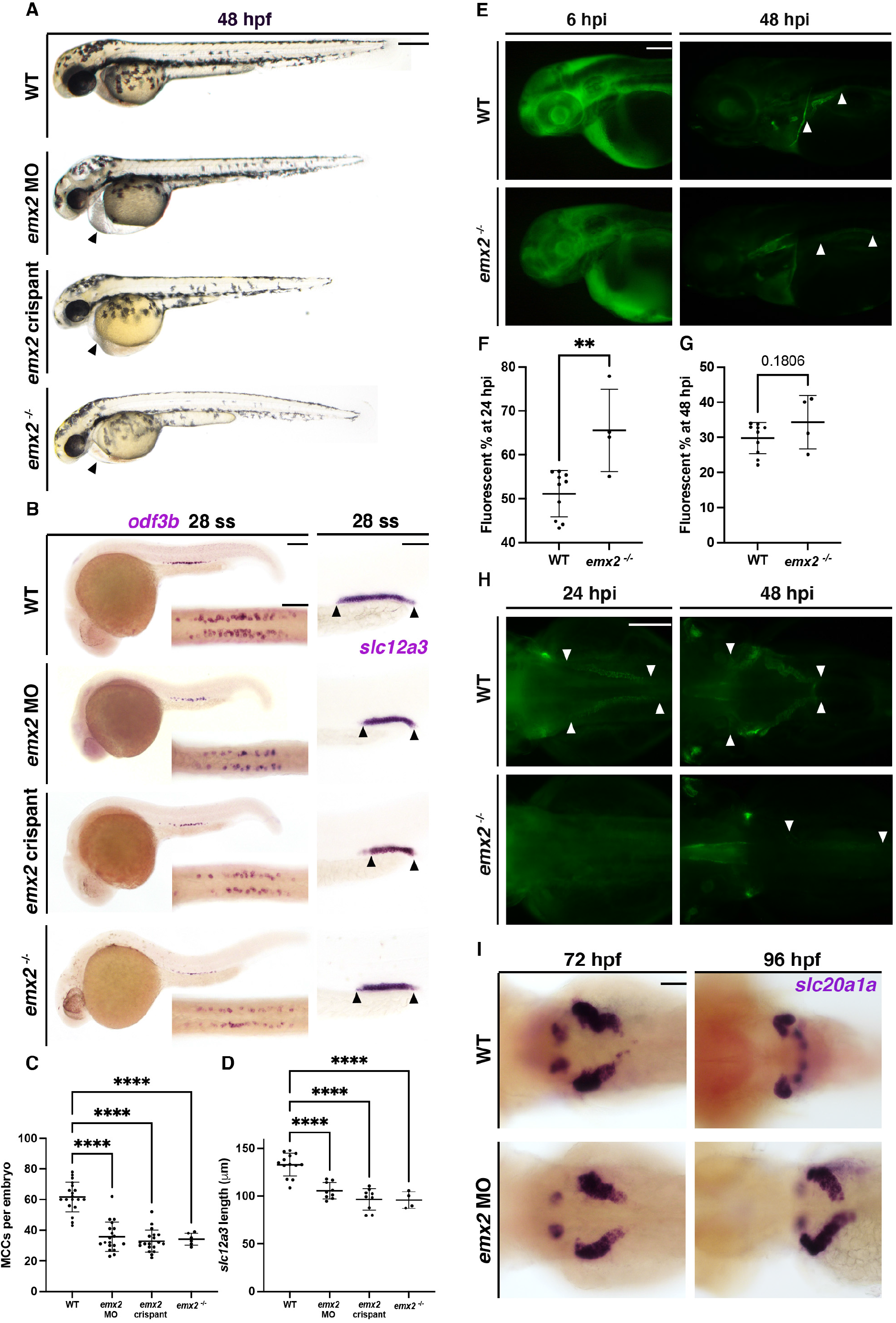
Loss of *emx2* causes phenotypes associated with cilia defects in zebrafish embryonic tubule. (A) WT, *emx2* MO, *emx2* crispant and *emx2^−/−^* embryos live imaging at 48 hpf. Scale bar = 200 μm. (B) 28 ss WT, *emx2* MO, *emx2* crispant and *emx2^−/−^* embryos stained via WISH using the MCC marker *odf3b* and the DL marker *slc12a3*. Scale bars = 100 μm, DL and MCC inset = 50 μm. (C) Number of MCCs per embryo between 28 ss WT, *emx2* MO, *emx2* crispant and *emx2^−/−^*embryos. (D) *slc12a3* length between 28 ss WT, *emx2* MO, *emx2* crispant and *emx2^−/−^* embryos. (E) Lateral view of WT and *emx2^−/−^* embryos at 6 hpi and 48 hpi after dextran-FITC injection. Scale bars = 100 μm. (F) Percentage of remaining fluorescent at 24 hpi and (G) percentage at 48 hpi as compared to 6 hpi between WT and *emx2^−/−^* embryos. (H) Dorsal view of WT and *emx2^−/−^*embryos at 24 hpi and 48 hpi after dextran-FITC injection. Scale bars = 100 μm. (I) 72 hpf and 96 hpf WT and *emx2* MO stained via WISH for *slc20a1a* to mark PCT. Scale bar = 50 μm. Data presented on graphs are represented as mean ± SD; * p<0.05, ** p< 0.01 ***p < 0.001 and ****p < 0.0001 (t-test or ANOVA).

Next, we wondered if this decrease in MCCs was associated with changes in MCC specification. MCC fate choice from the renal progenitor state occurs during the 10-26 ss, where MCC progenitors can be discerned based on expression of the transcription factor encoded by *pax2a* and components of the Notch signaling pathway like the ligand encoded by *jag2b* (Liu et al., 2007; Ma and Jiang, 2007; Marra et al., 2019). WISH analysis revealed a statistically significant reduction in the number of *pax2a^+^* and *jag2b^+^* cells in *emx2* deficient embryos compared to WT, respectively (Supplemental Figure 4B,C,E,F). These observations are consistent with a role for Emx2 in directing MCC fate choice.

Interestingly, using the marker *slc12a3*, we also found a significant decrease in the length of the DL segment in *emx2* deficient embryos compared to WT (Figure 2B,D). We also surveyed other nephron segments using WISH and found no difference between WT and *emx2* deficient embryos in the length of the PST segment based on expression of *trpm7* or the DE segment based on expression of *slc12a1* (Supplemental Figure 5A-D). In addition, to determine if the observed changes in nephron segments in the *emx2* deficient embryos were related to changes in embryo size, we performed several body measurements in WT and *emx2* deficient embryos. Neither the distance from the tip to tail, distance from yolk to cloaca, or length of the yolk extension differed between WT and *emx2* deficient embryos (Supplemental Figure 5E-H). Therefore, we concluded that the changes observed in DL and MCC numbers in *emx2* deficient embryos reflect direct roles of *emx2* in pronephros development as opposed to indirect consequences related to changes in the size of the embryo. These results led us to conclude that *emx2* is essential for renal lineage development in the zebrafish pronephros.

To further explore if the decreases in renal MCCs were functionally significant, we next assessed nephron clearance. We microinjected WT and *emx2*^−/−^ embryos with 40 kDa Dextran-FITC at 48 hpf, then performed live imaging at 6 hours post injection (hpi), 24 hpi and 48 hpi. Interestingly, we observed a delay in net fluid clearance in *emx2*^−/−^ embryos, as well as delayed coiling morphogenesis of the PCT segment at 24 hpi and 48 hpi (Figure 2E-H). As previous studies have shown that nephron fluid flow plays an important role in PCT morphogenesis and cell migration (Vasilyev et al., 2009), we performed WISH using the PCT marker *slc20a1a.* Similarly, we observed a delay in PCT coiling in *emx2* morphants compared to WT embryos at 72 hpf and 96 hpf (Figure 2I).

### *emx2* deficiency causes several body morphology defects

Defects in *emx2* are known to affect a plethora of tissues such as brain, ear hair cell development, kidney and reproductive tracts in mice and humans (Pellegrini et al., 1996; Miyamoto et al., 1997; Yoshida et al., 1997; Tole et al., 2000; Pang et al., 2008; Holley et al., 2010; Jiang et al., 2017; Li et al., 2022). In addition, our WISH data demonstrated a broad range of tissues labeled with *emx2* transcripts. Therefore, we also hypothesized that *emx2* deficiency in zebrafish will also cause several body morphology defects. Indeed, we observed pleiotropic phenotypes in *emx2* deficient embryos in addition to previously shown phenotypes in kidney and other ciliated organs. For instance, Alcian Blue staining at 96 hpf revealed a gaping jaw in *emx2* morphants compared to WT embryos. In contrast to WT embryos which formed five ceratobranchial (CB) bones, *emx2* morphant formed less CB bones (Supplemental Figure 6A). WISH staining using fin bud progenitor marker *mecom* showed a decreased fin bud area in *emx2* morphant compared to WT embryos (Supplemental Figure 6B,C). o-Dianisidine staining for erythrocytes at 48 hpf revealed the presence of blood clots around the head, eyeball and decrease in blood accumulation below the yolk ball in *emx2* morphants compared to WT controls (Supplemental Figure 6D). Furthermore, Acridine Orange staining at 24 hpf revealed increased cell death across the body in *emx2* morphant compared to WT embryos, such as the brain, somites and cloaca (Supplemental Figure 6E-I). Overall, these results indicate that *emx2* is involved in several physiological processes throughout the body, such as regulating cartilage and fin development, vasculature, brain, and kidney development, which are consistent with previous studies.

### *emx2* induces prostaglandin biosynthesis in renal cilia development

In the zebrafish embryo, decreased MCCs in conjunction with a reduced DL nephron segment are hallmark characteristics of renal cilia defects due to reduction in prostaglandin biosynthesis from alterations in the function of the transcription coactivator encoded by *ppargc1a* (Chambers et al., 2020). Interestingly, previous studies have shown that myoblasts overexpressed with EMX2 had a 7.64-fold increase in *ppargc1a* (PGC-1α in mammal) expression (Hatch et al., 2017). Given our observed phenotype with decreased DL length and MCC numbers in our *emx2* deficient embryos, we hypothesized that *emx2* is upstream of *ppargc1a* in regulating prostaglandin biosynthesis in renal cilia development. *ppargc1a* has recently been shown to regulate key players in prostaglandin biosynthesis. Importantly, deficiency of *ppargc1a* significantly reduces the quantity of *ptgs1* transcripts in the 24 hpf zebrafish embryo, where addition of *ptgs1* transcripts is sufficient to rescue cilia phenotypes in these *ppargc1a* deficient embryos (Chambers et al., 2020). Further, prostaglandin E_2_ (PGE_2_) is a molecule derived from PGH_2_, a prostaglandin precursor, and a major prostanoid detected in the zebrafish embryo (Cha et al., 2005). PGE_2_ is a product of biosynthesis reactions catalyzed by *ptgs1* and *ptgs2* (also known as COX1 and COX2 respectively) from arachidonic acid (Lord et al., 2007), and addition of PGE_2_ also restores cilia formation and renal MCC fate in *ppargc1a* deficient embryos (Chambers et al., 2020).

To study if *emx2* induces prostaglandin biosynthesis in renal cilia development, we first surveyed the promoter region of zebrafish *ppargc1a* and *ptgs1* for consensus sites for Emx2. Using the previously reported Emx2 consensus binding site (Hatch et al., 2017); we found 3 sites with striking similarities to the reported consensus site located within 10 kb upstream and downstream of *ppargc1a* open reading frame (ORF) for both strands (Supplemental Figure 7E). Similarly, we also found 4 sites within 10 kb upstream and downstream of the *ptgs1* ORF for both strands (Supplemental Figure 7F). The prevalence of consensus sites suggested potential interaction between Emx2 and these target genes. To investigate, we conducted WISH for *ppargc1a* at 28 ss. We observed a reduced domain of *ppargc1a* expression in the *emx2* morphant pronephros compared to WT embryos, and expression was qualitatively diminished elsewhere as well, such as the somites (Figure 3A,B). Additionally, we also observed a reduction of the *ppargc1a* pronephros domain in *emx2* morphant compared to WT embryos at earlier stages such as 15 ss or 20 ss (Supplemental Figure 7A-D). WISH studies also revealed a reduction in the pronephros domain of the cyclooxygenase enzyme *ptgs1* at 28 ss, which is a downstream target of *ppargc1a* (Figure 3D,E). Furthermore, qRT-PCR analysis revealed a statistically significant reduction in *ppargc1a* and *ptgs1* transcripts in *emx2* morphant compared to WT embryos at 28 ss (Figure 3C,F). Next, we explored if *emx2* influences the production of the prostaglandin biomolecule prostaglandin E_2_ (PGE_2_). We used a commercially available ELISA assay to measure endogenous PGE_2_ level between WT and *emx2* morphant embryos at 28 ss stage. There was no significant difference in the average amount of endogenous PGE_2_ in *emx2* morphants compared to WT embryos (Supplemental Figure 7G). This was surprising given the marked regional expression changes of *ppargc1a* and *ptgs1* observed in *emx2* deficient embryos, and therefore we hypothesize that there are important regional differences in endogenous PGE_2_ levels that affect the kidney which were not captured in the ELISA assay which relies on the entire embryo.

**Figure 3.**
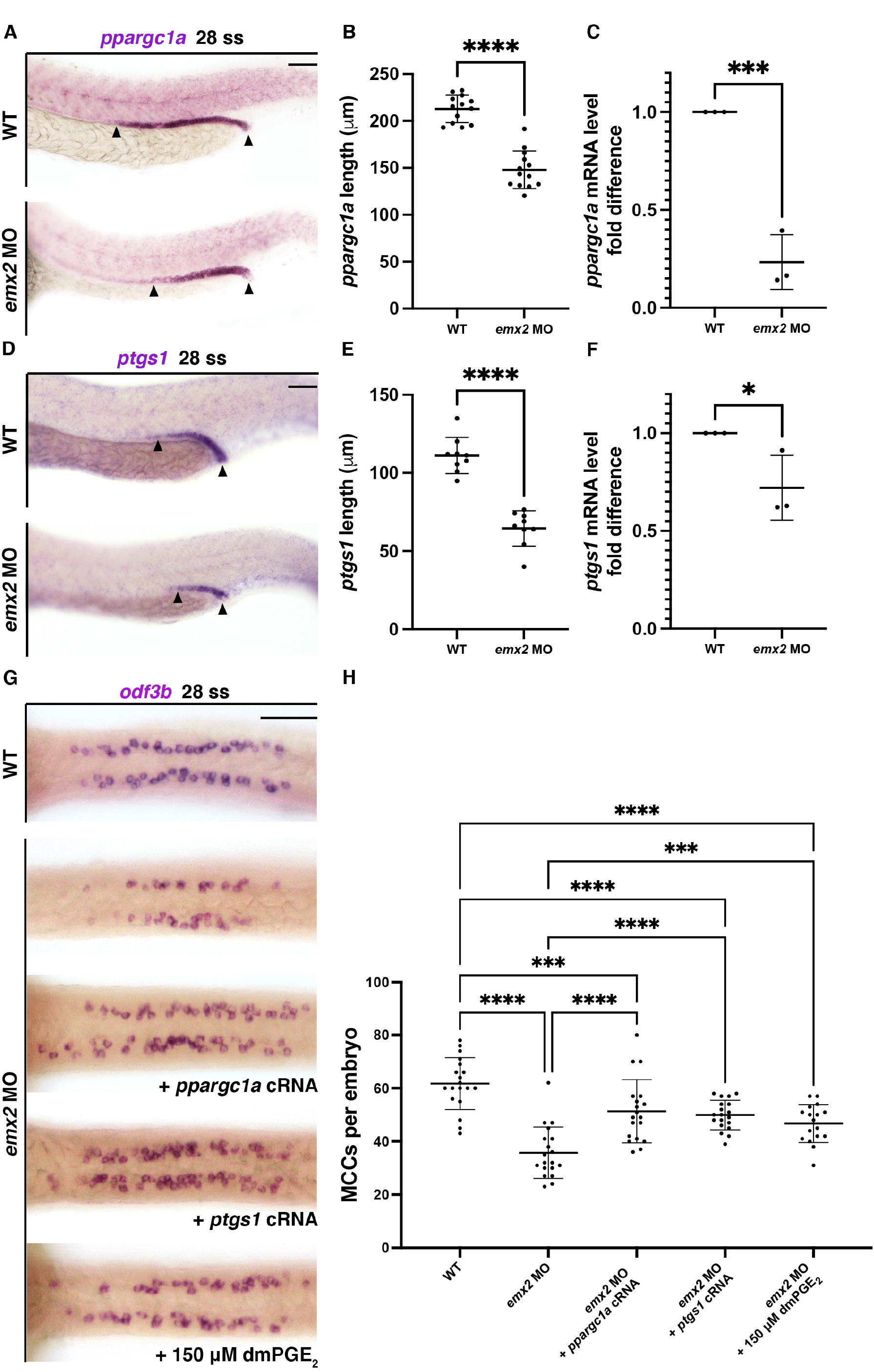
*emx2* induces prostaglandin biosynthesis in renal cilia development. (A) 28 ss WT and *emx2* MO embryos stained via WISH for *ppargc1a*. Scale bar = 50 μm. (B) *ppargc1a* length between 28 ss WT and *emx2* MO embryos. (C) Relative *ppargc1a* mRNA expression between WT and *emx2* MO embryos. (D) 28 ss WT and *emx2* MO embryos stained via WISH for *ptgs1.* Scale bar = 50 μm. (E) *ptgs1* length between 28 ss WT and *emx2* MO embryos. (F) Relative *ptgs1* mRNA expression in WT and *emx2* MO embryos. (G) 28 ss embryos stained via WISH for MCC marker *odf3b* for different treatment groups. Scale bar = 50 μm. (H) Number of MCCs per embryo between different treatment groups at 28 ss. Data presented on graphs are represented as mean ± SD; * p<0.05, ** p< 0.01 ***p < 0.001 and ****p < 0.0001 (t-test or ANOVA).

Given these observations, we sought to determine if the renal lineage defects in *emx2* deficient embryos could be rescued by *ppargc1a* transcripts, *ptgs1* transcripts or exogenous PGE_2_ treatment. Due to the similarities of phenotype between *emx2* morphant and *emx2^−/−^* embryos, and to reduce the number of animals used, here we performed rescue studies using *emx2* morphants due to the simplicity of sample generation. Interestingly, the provision of *ppargc1a* transcripts partially rescued MCC number in the *emx2* deficient embryos at the 28 ss (Figure 3G,H). Similarly, the provision of *ptgs1* transcripts partially rescued MCC number in *emx2* deficient embryos (Figure 3G,H). Additionally, to interrogate the effect of PGE_2_ in MCC development in *emx2* deficient embryos, we performed drug treatments in which we incubated *emx2* deficient embryos with dmPGE_2_, a more stable form of PGE_2_ that is amenable to chemical genetics, from 6 hpf until 24 hpf. *emx2* deficient embryos supplied with dmPGE_2_ had partially rescued MCC numbers per embryo compared to embryos treated with vehicle alone (Figure 3G,H). Altogether, these comprehensive results lead us to conclude that *emx2* promotes prostaglandin biosynthesis by controlling the expression of *ppargc1a, ptgs1* and ultimately affects local PGE_2_ production in the developing zebrafish.

### Deficiency of *emx2* leads to abnormal cilia formation and growth in renal tubule

Based on our data that *emx2* deficiency causes reduction in the number of MCCs per embryo and poor renal clearance, we hypothesized that *emx2* deficiency leads to abnormal cilia formation and growth in the epithelial cells of the nephron tubule. Cilia consist of a basal body that functions as a tubulin organizing center where the cilium grows from; a transition zone with proteins that help anchor cilia and function as a gate for proteins to travel between ciliary compartments; and lastly, an axoneme that protrudes from the cell and is comprised of microtubules that are covered by the unique ciliary membrane and ciliary tip (Fliegauf et al., 2007; Diener et al., 2015; Sun et al., 2019). To interrogate this hypothesis, we performed whole mount IF to detect acetylated α-tubulin (cilia), γ-tubulin (basal bodies) and DAPI (nucleus) in WT, *emx2* morphant and *emx2^−/−^* embryos at the 28 hpf stage (Figure 4A). We noticed that in the proximal pronephros, cilia length was significantly shorter in *emx2* morphant and *emx2^−/−^* embryos compared to WT controls (Figure 4B). We also observed a decrease in proximal fluorescent intensity of acetylated α-tubulin in *emx2* morphant and *emx2^−/−^* embryos compared to WT embryos (Figure 4C). Additionally, in the proximal pronephros, we observed a significant decrease in the percentage of ciliated basal bodies and the number of basal bodies and in both *emx2* morphant and *emx2^−/−^* embryos compared to WT (Figure 4D,E). Consistent with our observation that *emx2* deficiency leads to decreased MCC formation, our data suggest there is abnormal cilia formation and growth in the pronephros proximal tubule. To investigate our ciliopathic phenotype further, we examined the distal tubule via IF (Supplemental Figure 8A). Consistently, we also observed that cilia length in the distal tubule was significantly decreased in *emx2* morphant and *emx2^−/−^* embryos compared to WT (Supplemental Figure 8B). We also observed a decrease in the distal fluorescent intensity of acetylated α-tubulin in *emx2* morphant and *emx2^−/−^* embryos compared to WT embryos (Supplemental Figure 8C). Similarly, there was a decrease in percentage of ciliated basal bodies in *emx2* morphant and *emx2^−/−^* embryos compared to WT (Supplemental Figure 8D). *emx2* deficient embryos also exhibited fewer numbers of basal bodies compared to WT in the proximal pronephros (Figure 4E), and *emx2^−/−^* mutants similarly had decreased basal bodies in the distal pronephros (Supplemental Figure 8E). Altogether, our data indicated that cilia formation is disrupted in both the proximal and distal domain in *emx2* deficient embryos, and support our conclusion that *emx2* is essential for renal ciliogenesis.

**Figure 4.**
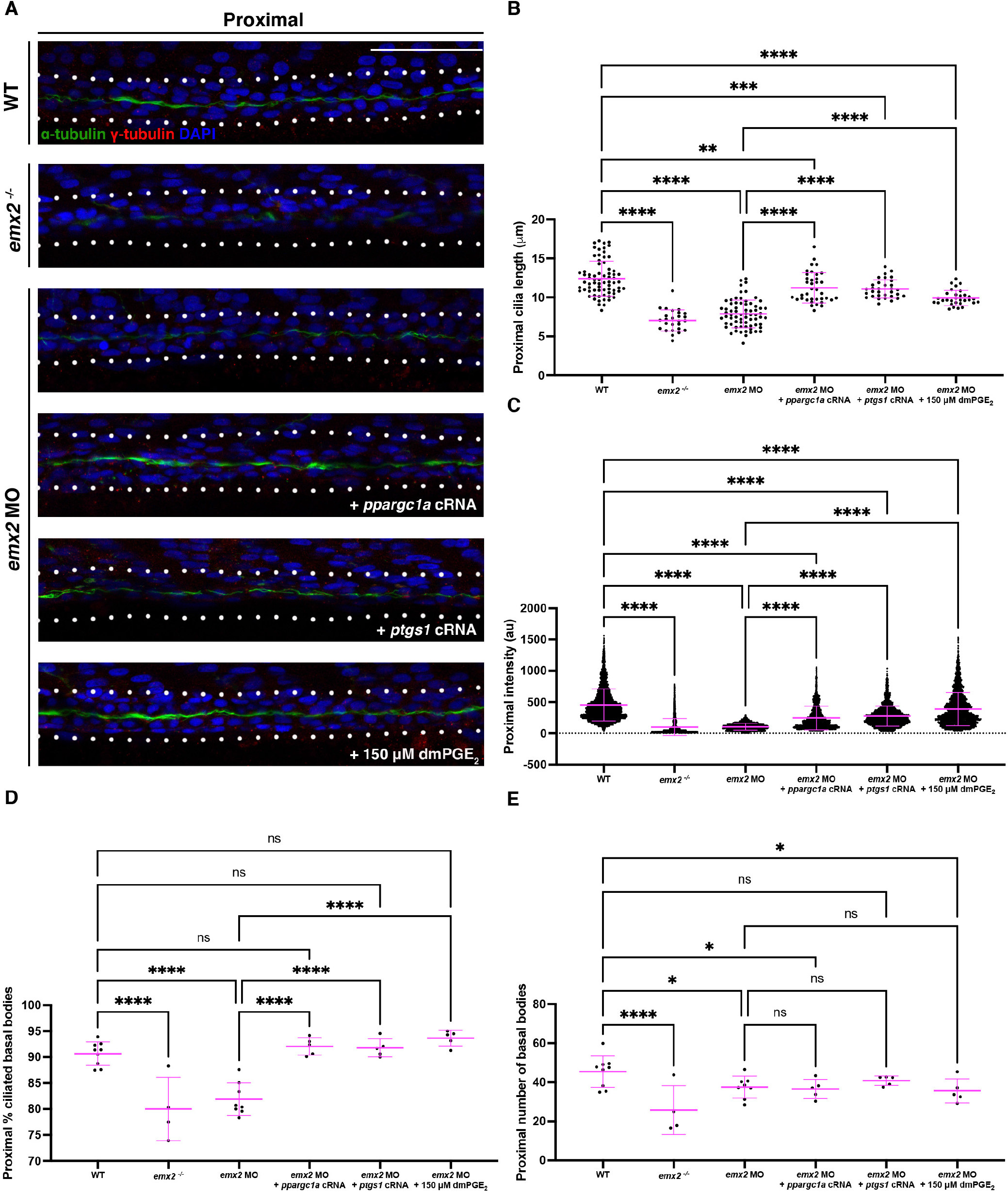
Cilia loss in proximal pronephros due to *emx2* deficiency can be rescued by supplement of key factors in prostaglandin biosynthesis. (A) 28 hpf whole-mount IF for acetylated α-tubulin (cilia, green), γ-tubulin (basal bodies, red), and DAPI (nucleus, blue) in the proximal pronephros between different treatment groups. Scale bar = 50 μm. (B) Cilia length for the proximal pronephros between different treatment groups. (C) Fluorescence intensity plot of α-tubulin intensity in the proximal pronephros between different treatment groups. (D) Percentage of ciliated basal bodies/total basal bodies in the proximal pronephros between different treatment groups. (E) Number of basal bodies in the proximal pronephros between different treatment groups. Data presented on graphs are represented as mean ± SD; * p<0.05, ** p< 0.01 ***p < 0.001 and ****p < 0.0001 (t-test or ANOVA).

### *emx2* mediated ciliogenesis operates via prostaglandin biosynthesis factors

Since the overexpression of prostaglandin biosynthesis factors (*ppargc1a, ptgs1)* or dmPGE_2_ treatment successfully rescued MCC formation in *emx2* morphants, next we hypothesized that the overexpression of prostaglandin biosynthesis factors would also rescue ciliogenesis within *emx2* morphants. Indeed, provision of *ppargc1a* transcripts to *emx2* deficient embryos rescued cilia length, acetylated α-tubulin intensity and percentage of ciliated basal bodies in both the proximal and distal domain of the pronephros (Figure 4A-D; Supplemental Figure 8A-D). Similarly, provision of *ptgs1* transcripts to *emx2* deficient embryos restored ciliogenesis throughout the nephron tubules, as did incubation with exogenous dmPGE_2_ (Figure 4A-D; Supplemental Figure 8A-D). Interestingly, *ppargc1a* transcripts, *ptgs1* transcripts or dmPGE_2_ did not rescue the average number of proximal or distal basal bodies in *emx2* morphants (Figure 4E, Supplemental Figure 8E). From this, we hypothesize that *emx2* has an essential role in basal body formation that does not operate through prostaglandin signaling. Collectively, these results enable us to discern that the ciliary growth defects in the pronephros from *emx2* loss of function resulted from disruptions in PGE_2_ biosynthesis, specifically establishing that *emx2* regulates the Ppargc1a/Ptgs1/PGE_2_ signaling cascade to control renal ciliogenesis.

### *emx2* regulates basal body development in renal cilia

The basal body is an important structure of the cilium. The basal body is made up of a centriole, which is the locus where cilia growth happens (Reese., 1965; Preble et al., 1999; Corkins et al., 2021). After the centriole is generated, basal bodies migrate and fuse with the apical surface of multiciliated cells. Basal body docking, thus, is an important and essential phenomenon for proper cilia development. Previous studies on *emx2* have demonstrated a significant role of Emx2 in controlling hair bundle polarity reversal in sensory hair cells in the ear hair cells and neuromasts, especially in positioning the basal body from which the sensory cilia grow from (Holley et al., 2010; Jiang et al., 2017; Ji et al., 2018; Jacobo et al., 2019; Kozak et al., 2020). However, there has not been much known about the role of *emx2* in basal body positioning in renal cilia. Due to an implicated role of *emx2* in basal body positioning, we hypothesized that *emx2* deficient embryos may have aberrant basal bodies compared to WTs.

To investigate basal body positioning in zebrafish renal cilia, we performed whole mount IF to detect acetylated α-tubulin (cilia), γ-tubulin (basal bodies) and DAPI (nucleus) in WT and *emx2* deficient embryos at the 28 hpf stage (Figure 5A). We examined whether cells that make up the pronephric tubule possessed basal bodies lacking proper apical orientation. Interestingly, we identified basal bodies with non-apical orientation in the proximal tubule in *emx2^−/−^* and *emx2* morphant embryos compared to WT embryos (Figure 5A). To quantify this, we scored each of those basal bodies as an aberrant basal body. There was a statistically significant increase in the incident of aberrant basal bodies per 100 μm in the proximal tubule in both *emx2* deficient groups compared to WT (Figure 5B). Next, we found that there was an increase in the percentage of individual cells that make up the proximal tubule with aberrant basal bodies in both *emx2* deficient groups compared to WT (Figure 5C). We also found an increase in the overall percentage of aberrant basal bodies in the proximal tubule in both *emx2* deficient groups compared to WT (Figure 5D). We also surveyed the distal tubule for aberrant basal bodies. Similar to the proximal tubule, there was an increased incidence of basal bodies with non-apical orientation in *emx2^−/−^* and *emx2* morphant embryos compared to WT embryos (Figure 5A). Interestingly, in both proximal and distal tubule, we observed misplaced basal bodies often with stunted cilia projections (Figure 5A). Similar to the proximal tubule, we observed an increase in the incident of aberrant basal bodies per 100 μm in the distal tubule in both *emx2* deficient groups compared to WT (Figure 5E). Additionally, we observed an increase in the percentage of DAPI cells that make up the distal tubule with aberrant basal bodies in both *emx2* deficient groups compared to WT (Figure 5F), and an increase in the percentage of aberrant basal bodies in the distal tubule in both *emx2* deficient groups compared to WT (Figure 5G). Our data show that the deficiency of *emx2* leads to incorrect basal body location in both the proximal and distal renal tubule cells. Thus, *emx2* serves an important role in basal body positioning within the zebrafish pronephros.

**Figure 5.**
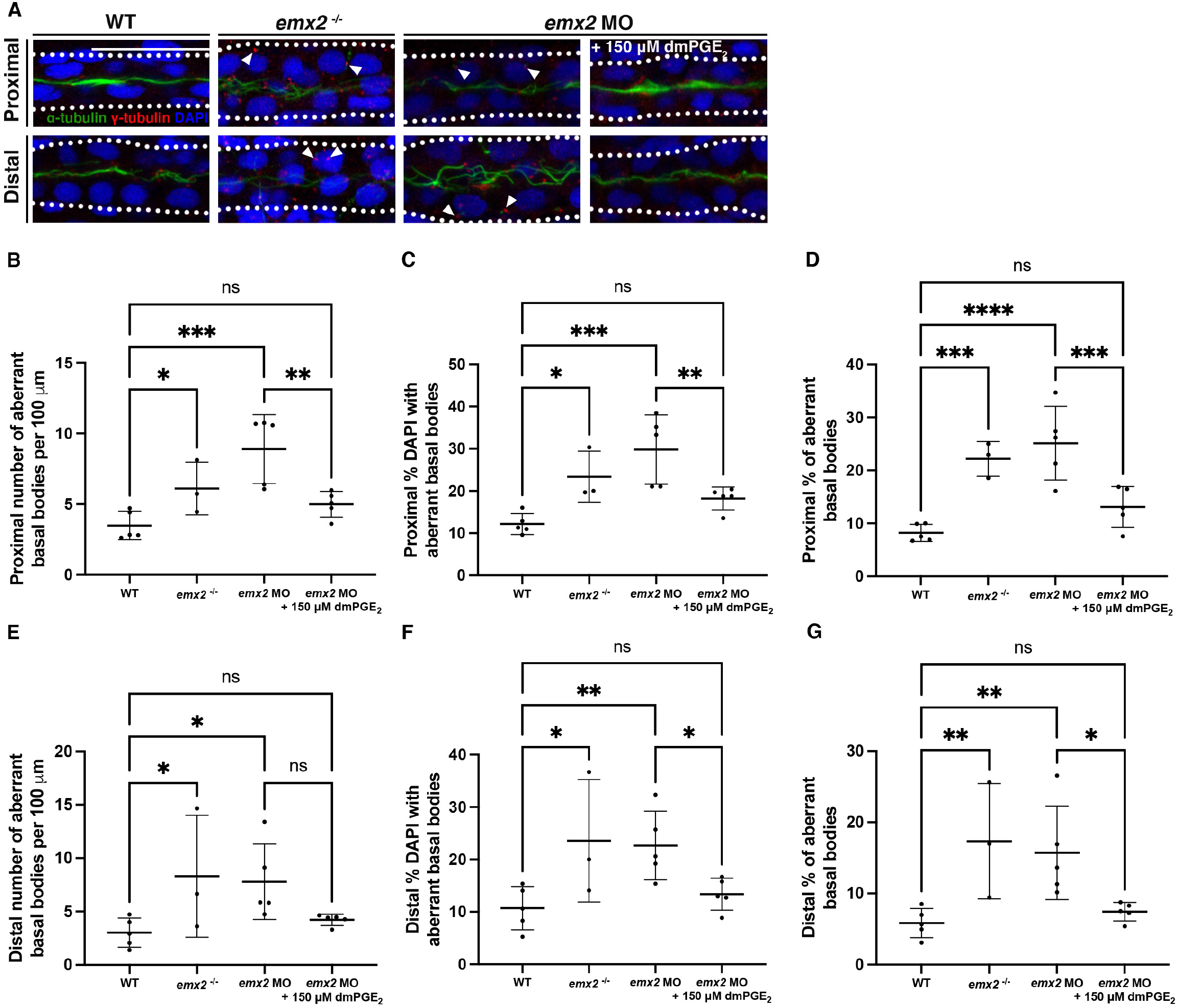
Basal body aberrancy in pronephric tubule due to *emx2* deficiency can be rescued by prostaglandin. (A) 28 hpf whole-mount IF for acetylated α-tubulin (cilia, green), γ-tubulin (basal bodies, red), and DAPI (nucleus, blue) in the proximal and distal pronephros between different treatment groups. White arrows indicated aberrant basal bodies. Scale bar = 50 μm. (B) Proximal number of aberrant basal bodies per 100 μm between different treatment groups. (C) Proximal percentage of DAPI cells that make up the pronephric tubule with aberrant basal bodies between different treatment groups. (D) Proximal percentage of aberrant basal bodies between different treatment groups. (E) Distal number of aberrant basal bodies per 100 μm between different treatment groups. (F) Distal percentage of DAPI cells that make up the pronephric tubule with aberrant basal bodies between different treatment groups. (G) Distal percentage of aberrant basal bodies between different treatment groups. Data presented on graphs are represented as mean ± SD; * p<0.05, ** p< 0.01 ***p < 0.001 and ****p < 0.0001 (t-test or ANOVA).

As our data have shown that treatment with dmPGE_2_ successfully rescued MCC formation in *emx2* morphants, we were next interested in whether dmPGE_2_ treatment can rescue our aberrant basal body phenotypes in *emx2* deficient embryos. To do so, we injected *emx2* MO at 1-cell stage, and performed drug treatment with injected embryos with dmPGE_2_ starting from 6 hpf until 28 hpf. Next, we performed whole mount IF to detect acetylated α-tubulin (cilia), γ-tubulin (basal bodies) and DAPI (nucleus) at 28 hpf stage (Figure 5A). Interestingly, in the proximal tubule, we saw a rescue in the number or aberrant basal bodies per 100 μm, percentage of cells with aberrant basal bodies, and percentage of aberrant basal bodies in *emx2* morphants supplemented with dmPGE_2_ compared to *emx2* morphants alone (Figure 5B-D). Similarly, in the distal tubule, we saw a rescue in the number of aberrant basal bodies per 100 μm, percentage of cells with aberrant basal bodies, and percentage of aberrant basal bodies in *emx2* morphants supplemented with dmPGE_2_ compared to vehicle (Figure 5E-G). Together, these data indicate that PGE_2_ is sufficient to rescue defective basal body positioning in *emx2* deficient embryos, revealing for the first time that *emx2* mediated prostaglandin signaling is crucial for basal body positioning in the zebrafish pronephros.

### Modulation of PGE_2_ level restores ciliogenesis defects in the KV and rescues heart looping

Finally, we were curious to explore whether prostaglandin signaling was a component of the ciliogenesis defect in other cell types of *emx2* deficient embryos. To examine this, we performed drug treatments in which we supplied *emx2^−/−^* embryos with dmPGE_2_ from 6 hpf until 55 hpf. Interestingly, we found that *emx2^−/−^* embryos supplied with dmPGE_2_ exhibited a statistically significant increase in cilia length in the KV (Figure 1J). Furthermore, dmPGE_2_ treated *emx2^−/−^* embryos underwent normal cardiac looping, with a complete restoration of the *situs solitus* phenotype (Figure 1J,K). We hypothesize that an optimal level of prostaglandin signaling is necessary to support ciliary function in the KV, and thereby establish normal heart randomization. These results suggest that Emx2 regulates the prostaglandin molecule PGE_2_ in controlling heart randomization, suggesting a novel role of Emx2 in controlling L/R patterning through prostaglandin signaling, and may have implications for the development other non-renal ciliated cells during embryogenesis.

## DISCUSSION

Knowledge about the molecular mechanisms governing ciliated cell development is critical for understanding organogenesis, as well as addressing a broad spectrum of birth defects and diseases that emerge from ciliary dysfunction. The present work identifies the Emx2 transcription factor as essential for cilia formation across multiple organs and a director of ciliated cell fate choice during the process of kidney lineage establishment in zebrafish. Furthermore, we determined that Emx2 has a key role in regulating the recently identified Ppargc1a/Ptgs1 genetic pathway, which produces the prostanoid signaling molecule PGE_2_ (Supplemental Figure 1A). Specifically, our work has revealed that Emx2 mediated production of PGE_2_ controls monociliated cell versus MCC fate choice during nephrogenesis in the embryonic zebrafish kidney, where it also drives normal ciliogenesis in these cell types, and that this pathway is relevant to ciliogenesis in the KV where defects lead to abnormal L/R patterning of the heart.

The Emx2/EMX2 transcription factor is well known for playing crucial roles during development, and there have been many fascinating discoveries about its roles in mechanosensory hair cell ontogeny. *Emx2* expression was first noted in olfactory placodes and later epithelium in mice, organs known to be highly ciliated (Simeone et al., 1992b), and subsequent work revealed that Emx2 had a vital role in ear hair cell development, where mouse mutants formed 60% fewer auditory hair cells and exhibited disruption of cochlear hair cell polarity, with polarity reversal across striola being damaged as well (Holley et al., 2010). Conditional knockout of *Emx2* in hair cells has been found similarly to display lack of line of polarity reversal in the otolith (Ji et al., 2022). Several landmark studies have elucidated the important role of *emx2/Emx2* in regulating hair cell orientation by regulated by core planar cell polarity in neuromasts (Jiang et al., 2017; Lozano-Ortega et al., 2018; Jacobo et al., 2019) as well as mouse maculae (Tona and Wu, 2020). Recent work has also revealed that EMX2 acts to polarize GPR156, a G-protein coupled receptor to signal through Gαi and trigger a 180 degree reversal in hair cell orientation (Kindt et al., 2021). However, the intriguing notion that Emx2 might possess functions in aspects of ciliated cell development in non-sensory cell types has not been explored until the present work.

Here, our work has revealed that Emx2 is required for ciliogenesis across several tissues of the zebrafish. In the absence of Emx2, we found that shortened or disorganized cilia form in the KV, neuromasts, otic vesicle, and the pronephros. Further, basal body positioning is defective in pronephros tubule cells in *emx2* deficient embryos, recapitulating the role of Emx2 in sensory hair cells. Roles for EMX2 in the kidney have been a challenge to study because *Emx2* knockout mice die postnatally within a few hours and exhibit catastrophic organ abnormalities, prominently an absence of the kidneys and other parts of the urogenital system (Pellegrini et al., 1996; Yoshida et al., 1997). Thus, this has precluded the analysis of EMX2 function in cell fate specification or differentiation. However, here using the zebrafish, we have demonstrated that *emx2* controls components of renal cell fate decisions and differentiation within pronephric nephrons, and that both processes involve Emx2 modulation of prostaglandin biosynthesis during early embryogenesis.

Prostaglandin biosynthesis has long been studied as an important physiological process controlling cilia development. A decade ago, it was found that PGE_2_ is important for regulating anterograde intraflagellar transport in ciliogenesis (Jin et al., 2014). Regulation of prostaglandin signaling has been recently identified as a potential treatment for the condition nephronophthisis (NPH), a rare renal ciliopathy (Garcia et al., 2022). Using *Nphp1* KO mouse, a major gene important for primary cilium and cellular junctions, it was found that prostaglandin molecule PGE_1_ and Taprenepag, a PGE_2_ receptor agonist alleviated ciliopathy phenotypes in the *Nphp1* KO mouse (Garcia et al., 2022). This highlights the importance of understanding how prostaglandin signaling is regulated in the context of ciliogenesis. Here, by revealing the role of Emx2 as an essential inducer of prostaglandin biosynthesis during renal ciliogenesis, we address an important knowledge gap that is relevant to the treatment of ciliopathies.

Within MCC development, several studies have demonstrated that prostaglandin signaling is essential for renal ciliated cell development, where *ppargc1a* is requisite to control prostaglandin biosynthesis (Marra et al., 2019a; Chambers et al., 2020). Interestingly, overexpression of EMX2 in cultured muscle cells led to their upregulation of PGC-1α (also known as *ppargc1a* in zebrafish) (Hatch et al., 2017). Until now, however, the relationship between *emx2* and *ppargc1a* has not been explored during kidney development or ciliogenesis. Herein, our work has revealed that *emx2* is upstream of *ppargc1a* and controls prostaglandin biosynthesis to modulate both ciliogenesis and the process of renal ciliated cell fate choice between that of a monociliated and MCC identity. MCCs are found in the zebrafish kidney, but only in the human fetal kidney (Katz and Morgan, 1984) as well as in renal pathological conditions (Duffy and Suzuki, 1968; Larsen and Ghadially, 1974; Eymael, et al., 2022). Therefore, knowledge of MCC development and what genes are responsible for MCC development has relevance for understanding the pathogenesis of numerous kidney diseases.

Future studies are needed to delineate the relationship of Emx2 with other genetic factors during kidney development. Interestingly, Emx1 has been suggested to compensate for Emx2 deficiency in the telencephalon (Yoshida et al., 1997). Additionally, EMX1 and EMX2 expression have been found to overlap in the olfactory bulb and amygdaloid complex (Mallamaci et al., 1998). Given that *emx1* and *emx2* are paralogs of each other, and previous study has demonstrated the importance of *emx1* in nephron development in zebrafish (Morales et al., 2018), it is possible that *emx2* also interacts with *emx1* in kidney tubule development. Previous studies have shown that Notch1a downregulates Emx2 expression in controlling hair cell polarity (Jacobo et al., 2019; Kozak et al., 2020). Interestingly, previous studies have shown that Notch signaling is important for restricting MCC formation (Li et al., 2014; Marra and Wingert, 2016). Notch signaling has also been found to inhibit *etv5a*, a transcription factor essential for MCC development (Marra and Wingert, 2016). Therefore, there remains a question of how *emx2* interacts with Notch signaling in regulating MCC development in the zebrafish kidney.

Additionally, previous studies have shown a link between *Emx2* and WNT/β-catenin signaling in nephron development. qRT-PCR have demonstrated a decrease in *Emx2* and *Lim1* mRNA in β-catenin mutants when compared to WT. Additionally, β-catenin mutants also showed decreased expression of *Pax2*, *c-ret*, *Wnt11* and *Gdnf* compared to WT (Bridgewater et al., 2008). Notably, the knockdown of EMX2 has also been reported to promote canonical WNT signaling in lung cancer cells (Okamoto et al., 2010). Furthermore, previous studies found that *Pax2* regulates *Emx2* in the Wolffian duct in mouse, and *Emx2* participates in a complex renal gene regulatory network, including *Pax2/8*, *Gata3* and *Lim1* (Boualia et al., 2011; Boualia et al., 2013). Interestingly, most of these studies have been conducted in mouse models. Therefore, there remains open a question of whether mammalian *Emx2/EMX2* also participates in these physiological pathways as revealed here in the zebrafish kidney. Notably, given our understanding of the role of *emx2* in controlling MCC development in zebrafish kidney via prostaglandin biosynthesis, it becomes even more interesting to see how *emx2* and prostaglandin biosynthesis fits in this greater gene regulatory network in renal development.

Overall, the present study has further expanded our knowledge of Emx2 as an essential transcription factor across multiple developing organs. This work has revealed that Emx2 modulates prostaglandin biosynthesis to control renal ciliogenesis and kidney cell fate decisions, and further, that Emx2 control of ciliogenesis in non-renal cell types can also involve modulating prostaglandin signaling. These findings have broad implications to understanding the mechanisms of ciliated cell development and the basis of ciliopathic conditions.

Cox1: cyclooxygenase 1
Cox2: cyclooxygenase 2
dpf: days post fertilization
DE: distal early
DL: distal late
*emx2*: *empty spiracles homeobox gene 2*
IF: immunofluorescence
dmPGE_2_: 16,16-dimethyl-PGE_2_
hpi: hours post injection
IFT: intraflagellar transport
hpf: hours post fertilization
KV: Kupffer’s vesicle
MO: morpholino oligonucleotide
MCC: multiciliated cell
*ppargc1a*: *peroxisome proliferator-activated receptor gamma 1 alpha*
PKD: polycystic kidney disease
PGE_2_: prostaglandin E_2_
*ptgs1*: *prostaglandin-endoperoxide synthase 1*
PCT: proximal convoluted tubule
PST: proximal straight tubule
ss: somite stage
WISH: whole mount *in situ* hybridization
WT: wild-type

## METHODS

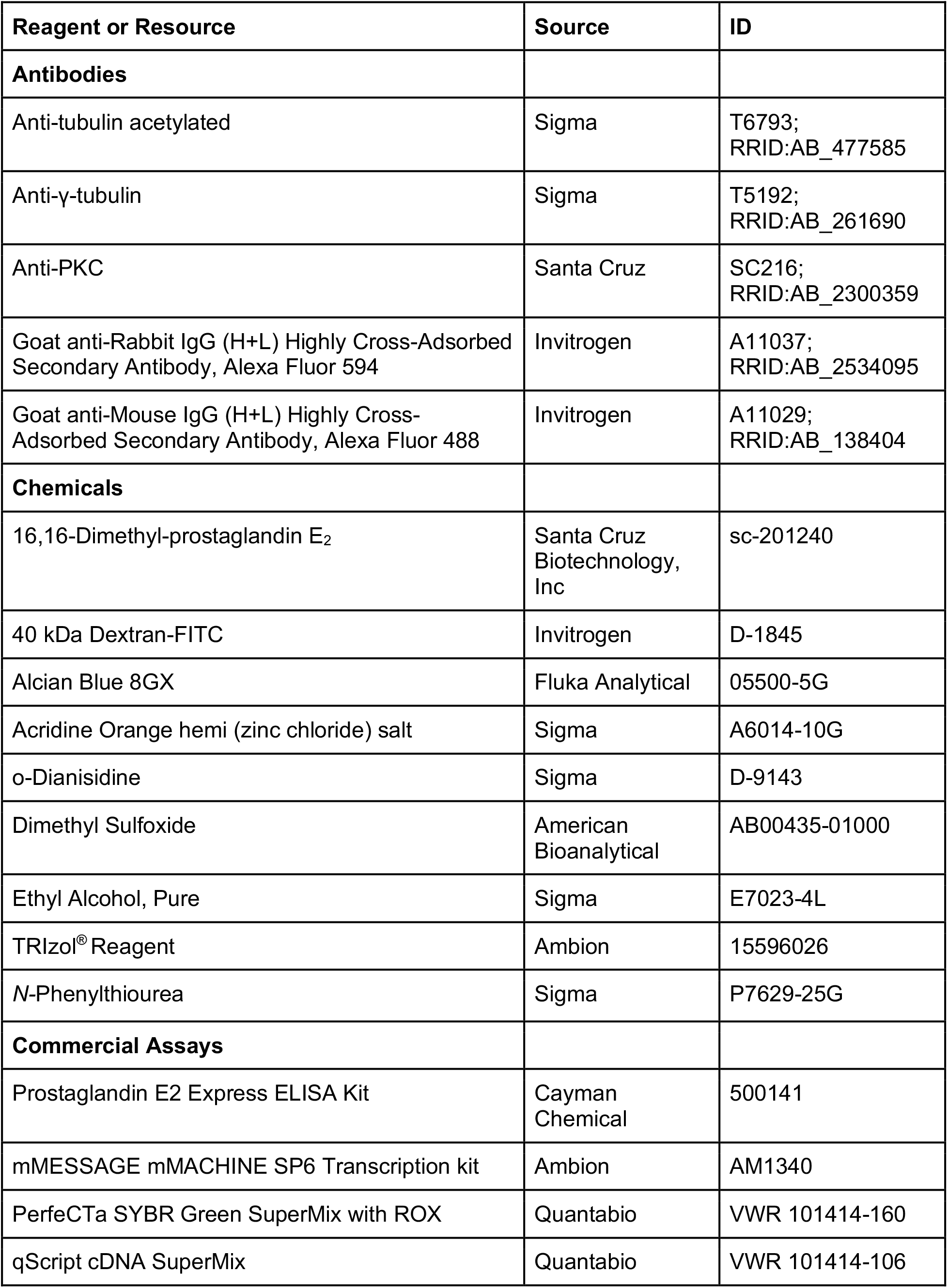
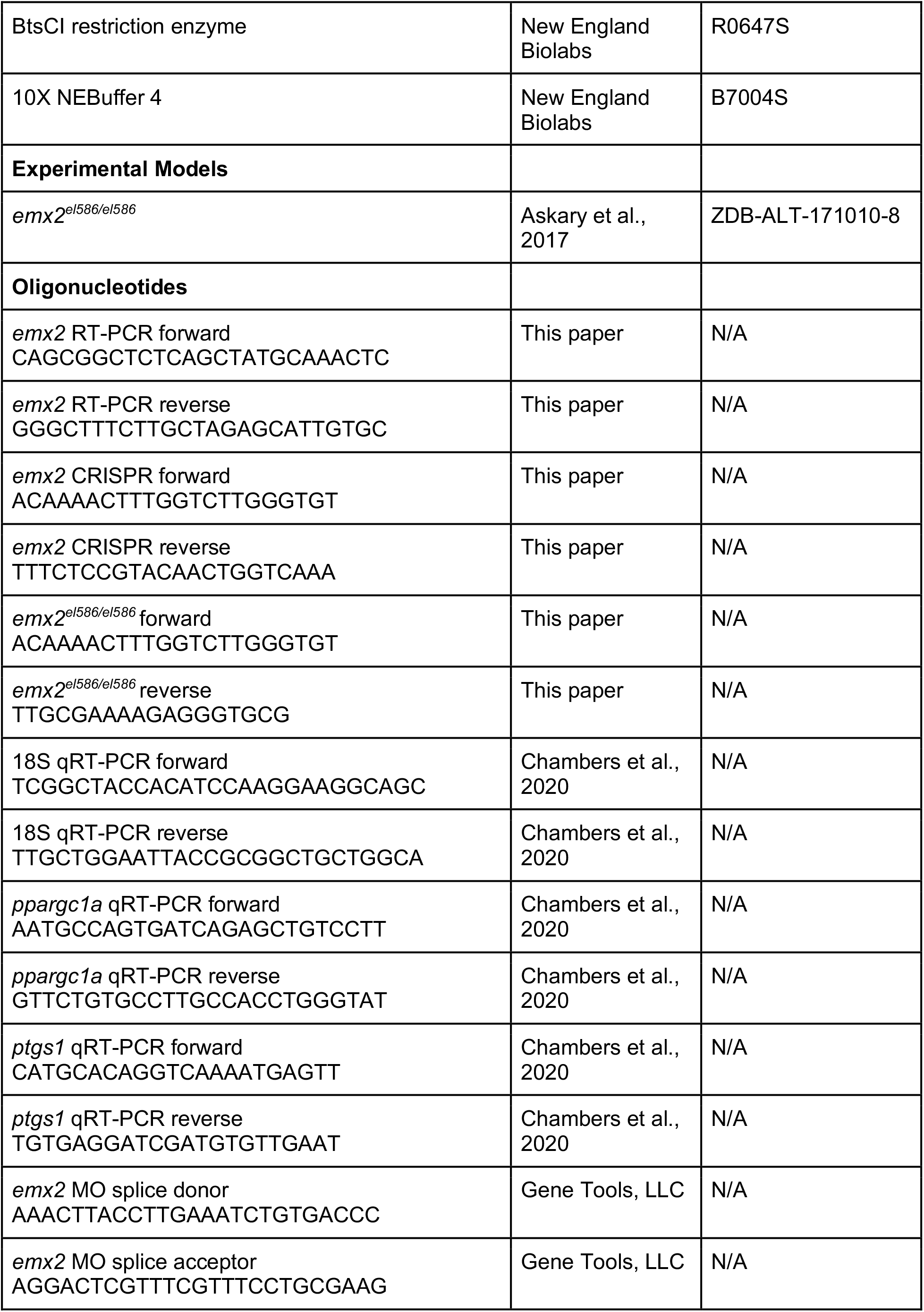
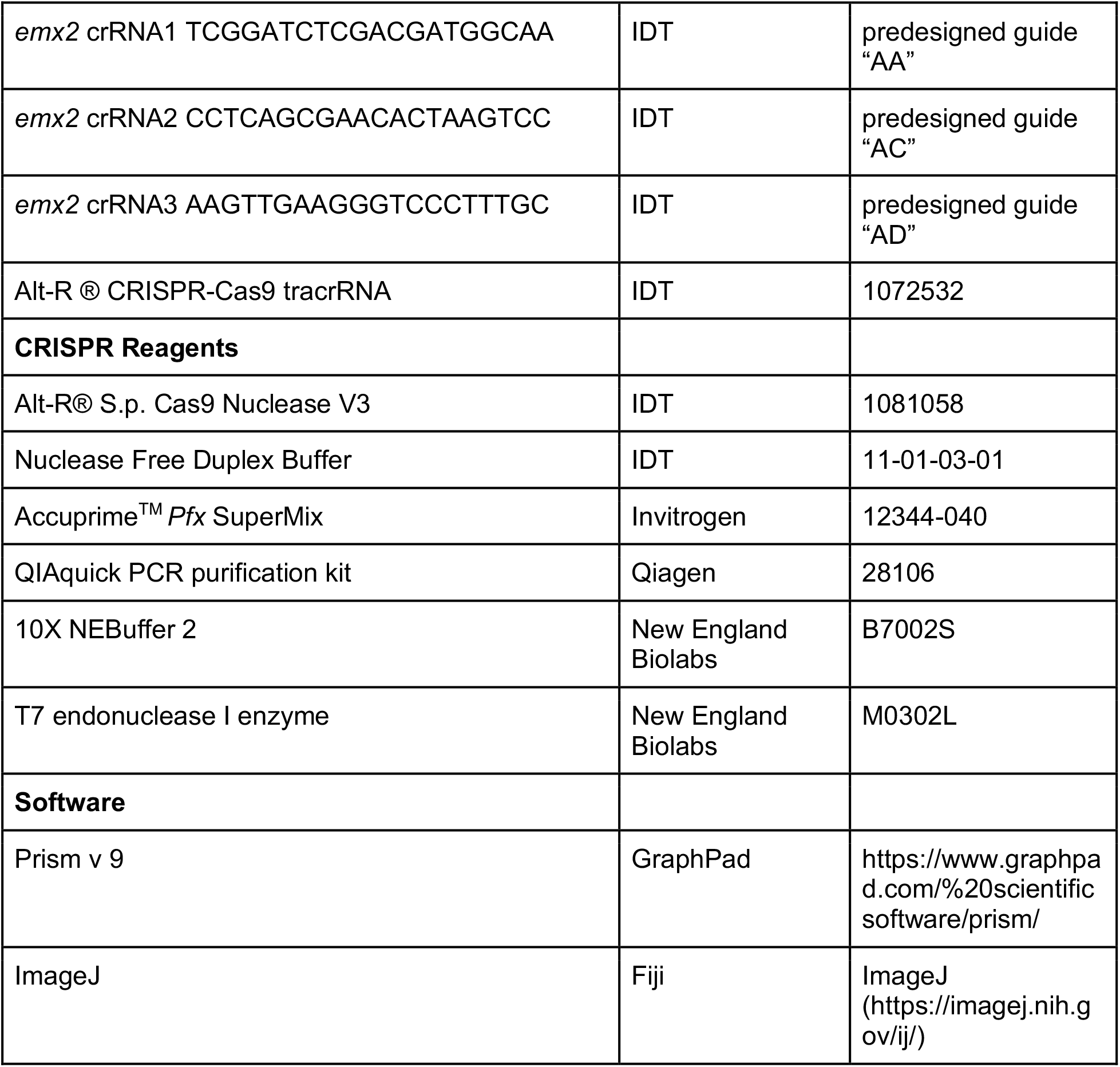

### EXPERIMENTAL MODEL AND SUBJECT DETAILS

The Center for Zebrafish Research at the University of Notre Dame maintained the zebrafish used in these studies and experiments were performed with approval of the University of Notre Dame Institutional Animal Care and Use Committee (IACUC), under protocol number 19-06-5412.

#### Animal models

Tübingen strain WT zebrafish were used for the reported studies. Zebrafish were raised and staged as previously described (Kimmel et al., 1995). For all experiments, embryos were incubated in E3 medium at 28°C until the desired developmental stage, anesthetized with 0.02% tricaine, and then fixed using either 4% paraformaldehyde/1x PBS (PFA), or Dent’s solution (80% methanol, 20% DMSO) (Westerfield, 2007). Embryos were analyzed before sex determination, so we cannot report the effect of sex and gender in the context of this study.

### METHOD DETAILS

#### Whole mount *in situ* hybridization (WISH)

WISH was performed as previously described (Cheng et al., 2014; Galloway et al., 2008; Lengerke et al., 2011). Linearized plasmids were transcribed in vitro with T7, T3 or SP6 enzymes to create antisense RNA probes either digoxigenin-labeled (*emx2, odf3b, myl7, slc12a3, slc20a1a, ptgs1, pax2a, jag2b, cetn4, trpm7, slc12a1, cdh17, mecom*) or fluorescein-labeled (*emx2, ppargc1a*) via *in vitro* transcription using IMAGE clone templates as previously described (Wingert et al., 2007; O’Brien et al., 2011; Gerlach and Wingert, 2014). All WISH experiments were performed in triplicate with a sample size of greater than 30 embryos for each replicate. Representative samples were imaged and analyzed.

#### Dextran-FITC injections

WT and *emx2^el586/el586^* heterozygous incrosses were incubated in 0.003% *N-*Phenylthiourea (Sigma). At 48 hpf, embryos were anesthetized and 40 kDa Dextran FITC conjugate (5 mg/mL) was injected into the somite to introduce to circulation (McCampbell et al., 2014; Kroeger et al., 2017). Embryos were imaged at 6 hpi, 24 hpi and 48 hpi. Mean fluorescent intensity of the head and the pericardium was calculated via ImageJ, and the percent of remaining fluorescence was calculated using the 6 hpi fluorescent intensity as a baseline.

#### Acridine Orange (AO) assay

AO experiments were conducted as described (Tucker and Lardelli, 2007; Kroeger et al., 2017). AO (Sigma) was prepared by dissolving 50 mg in 50 mL of MilliQ water to create a 100X solution and stored at −20°C, protected from light. AO was diluted in E3 to a 1X working solution before applied to 24 hpf live embryos. Embryos were incubated in AO/E3 for 30 minutes at room temperature, then washed 3 times for 10 min with E3. Samples were anesthetized in tricaine and imaged in methylcellulose.

#### Alcian Blue stain

Alcian Blue staining was conducted as described (Neuhauss et al., 1996). At 4 dpf, embryos were fixed in 4% PFA overnight at 4°C in glass vials. They were dehydrated in 100% MeOH at −20°C, then rehydrated. Embryos were bleached at room temperature for 1 h, then rinsed in PBST twice before digested in 1X proteinase K (10 mg/mL) for 15 minutes, then rinsed twice in PBST again. Afterward, embryos were stained overnight in 0.1% Alcian Blue dissolved in 70% ethanol/5% concentrated HCl in glass vials on the rocker. Embryos were then destained using acidic ethanol for 4 hours on the rocker, then rinsed twice in PBST and then rehydrated in ethanol series. Embryos were then stored in 100% glycerol for imaging.

#### Morpholino knockdown

Morpholino oligonucleotides (MO) were designed and obtained from Gene Tools, LLC. MOs were solubilized in DNase/RNase free water to generate 4mM stock solution, which were stored at −20°C. Embryos were injected at one-cell stage with 5 nl of diluted MO. The *emx2* MOs include an *emx2* splice donor 5’-AAACTTACCTTGAAATCTGTGACCC-3’ and *emx2* splice acceptor 5’-AGGACTCGTTTCGTTTCCTGCGAAG-3’. MO specificity was assessed using the reverse transcriptase polymerase chain reaction (RT-PCR) with the following pair of primers 5’-CAGCGGCTCTCAGCTATGCAAACTC-3’ and 5’-GGGCTTTCTTGCTAGAGCATTGTGC-3’. Products were isolated by PCR purification and Sanger Sequencing.

#### o-Dianisidine stain

o-Dianisidine powder (Sigma) was diluted in 100% ethanol and stored in 4°C. The working solution was prepared as follows: 2 ml of o-Dianisidine solution + 500 μl Sodium Acetate, pH = 4.5 + 2 ml distilled water + 100 μl 30 % H_2_O_2_. The working solution was applied to live embryos at 48 hpf in the dark at room temperature. Then, the reaction was stopped by washing 3X in distilled water, and embryos were then fixed in 4% PFA for imaging.

#### Immunofluorescence (IF)

Whole-mount IF experiments were completed as previously described (Gerlach and Wingert, 2014; Marra et al., 2017, 2019a; Chambers et al., 2020; Wesselman et al., 2023). For cilia, anti-tubulin acetylated diluted 1:400 (Sigma) was used. For basal bodies, anti-γ-tubulin diluted 1:400 (Sigma) was used. For apical membrane, anti-aPKC diluted 1:500 (Santa Cruz) was used.

#### PGE_2_ metabolite quantification

PGE_2_ metabolite quantifications were performed according to the manufacturer’s protocol (Cayman Chemical, Item No. 500141) and adapted from previously reported studies (Chambers et al., 2020; Wesselman et al., 2023). Groups of 25 WT and *emx2* MO embryos were pooled, anesthetized and flash frozen in 100% ethanol. Lysates were homogenized and the supernatant was isolated after centrifugation at 4°C (15000 rpm for 20 minutes). The kit reagents and manufacturer supplied protocol was followed for assay completion using a plate reader (SpectraMax ABSPlus) at 420 nm.

#### cRNA synthesis, microinjections and rescue studies

The *ppargc1a* ORF was cloned into a pUC57 vector flanked by a 5’ KOZAK sequence, Cla1 restriction site and a SP6 promoter region. On the 3’ side, the ORF was followed by a series of STOP codons, an SV40 poly A tail, a NotI restriction site and a T7 promoter region. *ppargc1a* RNA was generated by linearizing with Not1. Linearized plasmid was run off with SP6 RNA polymerase using the mMESSAGE mMACHINE SP6 Transcription kit (Ambion). *ppargc1a* RNA was injected together with *emx2* MO at one-cell stage at concentration 375 pg for rescue study. The *ptgs1* ORF was cloned into a pUC57 vector flanked by a 5’ KOZAK sequence, Cla1 restriction site and a SP6 promoter region. On the 3’ side, the ORF was followed by a series of STOP codons, an SV40 poly A tail, a NotI restriction site and a T7 promoter region. *ptgs1* RNA was generated by linearizing with Not1. Linearized plasmid was run off with SP6 RNA polymerase using the mMESSAGE mMACHINE SP6 Transcription kit (Ambion). *ptgs1* RNA was injected together with *emx2* MO at one-cell stage at concentration 825 pg for rescue study.

#### Rescue experiments with dmPGE_2_

Chemical treatments were adapted from previously described studies (Marra et al., 2019a; Chambers et al., 2020; Wesselman et al., 2023). In short, 16,16-Dimethyl-prostaglandin E_2_ (Santa Cruz Biotechnology) was dissolved in 100% DMSO (American Bioanalytical) to create 1 M stock. Then 1 M stock was diluted to the treatment dose. Treatments were performed in triplicate with n > 30 embryos per replicate.

#### qRT-PCR

Groups of 30 zebrafish embryos (WT and *emx2* morphants) were pooled at 24 hpf. RNA was extracted using Trizol (Ambion), and cDNA was made from RNA using qScript cDNA SuperMix (QuantaBio). The optimal cDNA concentration was 100 ng for *ppargc1a,* 100 ng for *ptgs1* and 1 ng for 18S controls. PerfeCTa SYBR Green SuperMix with ROX (QuantaBio) was used to complete qRT-PCR. The AB StepOnePlus qRT-PCR machine was used with the following program: 2 min 50°C hold, 10 min 95°C hold, then 35 cycles of 15 s at 95°C (denaturing) and 1 min at 60°C (primer annealing) and product extension steps. Each target and source were completed in three biological replicates and three technical replicates each, with the Ct value normalized to control. Data analysis was performed by using delta delta Ct values comparing WT embryos to the *emx2* morphant embryos, using 18S as a reference gene. The primers used in this study are: *ppargc1a* forward 5’-AATGCCAGTGATCAGAGCTGTCCTT-3’, *ppargc1a* reverse 5’-GTTCTGTGCCTTGCCACCTGGGTAT-3’, *ptgs1* forward 5’-CATGCACAGGTCAAAATGAGTT-3’, *ptgs1* reverse 5’-TGTGAGGATCGATGTGTTGAAT-3’, 18S forward 5’-TCGGCTACCACATCCAAGGAAGGCAGC-3’, 18S reverse 5’-TTGCTGGAATTACCGCGGCTGCTGGCA-3’.

#### CRISPR-Cas9 mutagenesis

We adapted methods from Hoshijima et al., 2019. gRNAs targeting the *emx2* coding region were selected using the pre-designed gRNAs from Integrated DNA Technologies (IDT). *emx2* sgRNA 1, *emx2* sgRNA 2, and *emx2* sgRNA 3 targeted different regions in exon 1 of the *emx2* coding frame. Selected crRNA and tracrRNA tools were obtained from IDT and dissolved into 100 μM IDT duplex buffer to make 100 μM stock. To form the crRNA:tracrRNA duplex, we combined a similar amount of cRNA and tracrRNA, vortexed and mixed and performed a rapid heat/cool program using a thermocycler to generate 50 μM duplex solution. To prepare 25 μM Cas9 stock solution, we obtained Cas9 protein from IDT and adjusted to 25 μM stock solution in 20 mM HEPES-NaOH (pH =7.5), 350 mM KCl, 20% glycerol and dispensed in aliquots and stored at −80°C. To prepare 5 μM cRNA:tracrRNA:Cas9 RNP complex, the crRNA:tracrRNA duplex was first diluted 1:1 with IDT duplex buffer. Then the injection mixes was prepared as follow: 1 μl 25 μM crRNA:tracrRNA (crRNA 1) + 1 μl 25 μM crRNA:tracrRNA (crRNA 2)+1 μl 25 μM crRNA:tracrRNA (crRNA 2) + 1 μl 25 μM Cas9 protein+2 μl nuclease-free water. The injection mix was incubated at 37°C for 5 minutes, then injected to the one-cell stage embryos with 3 nl. Crispants were identified by performing PCR with the following primers flanking the mutation site: forward 5’-ACAAAACTTTGGTCTTGGGTGT-3’ and reverse 5’-TTTCTCCGTACAACTGGTCAAA-3’. Primers were designed using CHOPCHOP web-based tool (https://chopchop.cbu.uib.no/) to flank all three sgRNA target sites in exon 1, respectively (Supplemental Figure 3). In brief, DNA was prepared from individual animals and Accuprime^TM^ *Pfx* SuperMix (Invitrogen) was used to amplify target sites. PCR products were purified using the QIAquick PCR purification kit (Qiagen) and sent to Notre Dame Genomics Core for sequencing analysis. Additionally, T7 endonuclease assay was used to confirm genome editing, adapted from the method described in Chambers et al., 2020. Purified PCR product (∼300 ng), 2 μl 10X NEBuffer 2 (New England Biolabs) and nuclease-free water (total of 20 μl) were mixed and rehybridized using the following program: 5 min at 95°C, ramp down to 85°C at 2°C/s, ramp down to 25°C at 0.1°C/s, and 25°C for 10 min. Rehybridized products were incubated with 1.5 μl T7 endonuclease I enzyme (New England Biolabs) at 37°C for 1 h and separated on 1.5% agarose gel.

#### Genetic models: *emx2*^el586/el586^ mutant line

The *emx2*^el586/el586^ mutant line was obtained from Lindsey Barske (Cincinnati Children’s Hospital, Ohio) (Askary et al., 2017). Mutant embryos and heterozygous adults were identified by performing PCR with the following primers flanking the mutation site: forward 5’-ACAAAACTTTGGTCTTGGGTGT-3’, and reverse 5’-TTGCGAAAAGAGGGTGCG-3’. PCR products were purified using the QIAquick PCR purification kit (Qiagen) and sent to Notre Dame Genomics Core for sequencing analysis. Additionally, restriction enzyme digest assay with BtsCI (New England Biolabs) was performed to identify WT, heterozygous or mutant samples. The PCR products (∼300 ng) were incubated with 2 μl 10X NEBuffer 4 (New England Biolabs), 1 μl BtsCI restriction enzyme, and nuclease-free water for a total of 20 μl. The mixture was incubated at 50°C for 1 hour, and separated on 2% agarose gel.

#### Image acquisition and phenotype quantification

A Nikon Eclipse Ni with a DS-Fi2 camera was utilized to take pictures of WISH samples and live imaging of zebrafish. WISH segment length was quantified using the Nikon Elements software polyline tool. To mount live embryos, methylcellulose with trace amounts of tricaine was used. IF images were acquired using a Nikon A1R confocal microscope.

#### Quantification and statistical analysis

ImageJ/Fiji software tool was used to quantify cilia phenotypes. Measurements were performed on samples imaged at 60X magnification. Counting was performed using the multi-point tool. Segment line tool was used for measuring lengths. Measuring fluorescent intensity was performed using the plot profile function. Each experiment was completed in a minimum of triplicate. From measurements, an average and standard deviation (s.d.) were calculated, and either one-way ANOVA or unpaired t-tests were completed to compare between WT and experimental groups via GraphPad Prism 9 software. Statistical details for each experiment are located in the figure legends.

## ACKNOWLEDGEMENTS

We thank the staff of the Department of Biological Sciences for support and the Center for Zebrafish Research at the University of Notre Dame for exceptional care of our zebrafish system. We are grateful for the members of Wingert Lab for assistance with this work. This work was supported by funds to R.A.W. from the College of Science and the generosity of the Gallagher Family for their support of stem cell research. We thank the Notre Dame Integrated Imaging Facility for providing us with the imaging facility that allowed us to take several images from this manuscript, and especially Sara Cole for her incredible expertise and guidance. We would like to thank Dr. Lindsey Barske for generously providing us with the *emx2^el586/el586^* zebrafish line. T.K.N would like to personally thank R.A.W, R.R and P.B for incredible mentorship.

## SUPPLEMENTAL FIGURE LEGENDS

**Supplemental 1.**
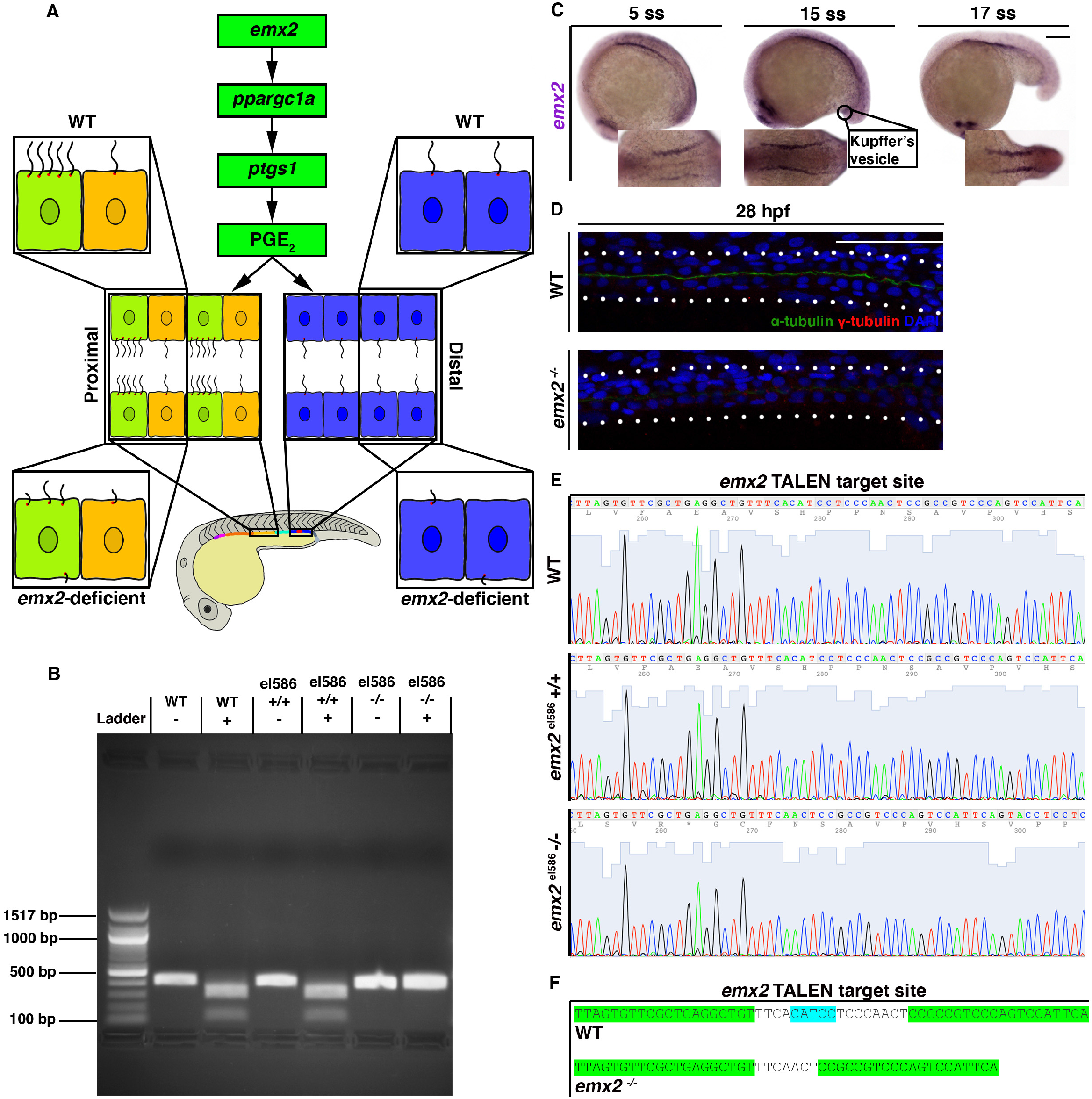
(A) Working model showing *emx2* induces prostaglandin biosynthesis for cilia development in zebrafish pronephros in both proximal and distal domain. (B) Image of DNA products after verifying with BtsCI restriction enzyme assay. − denotes samples without BtsCI and + denotes samples with BtsCI. WT embryos and confirmed WT embryos from *emx2^el586/el586^* line show two bands at 302 bp and 103 bp, while confirmed *emx2^−/−^* embryos show one band at 405 bp. (C) WT embryos stained via WISH using marker for *emx2* from 5 ss to 17 ss. Scale bar = 100 μm. (D) 28 hpf whole-mount IF for acetylated α-tubulin (cilia, green), γ-tubulin (basal bodies, red), and DAPI (nucleus, blue) in the distal pronephros of WT and *emx2^−/−^* embryos. Scale bar = 50 μm. (E) Sanger sequencing between WT, WT from *emx2^el586/el586^* line and *emx2^−/−^* embryos. (F) TALEN target site between WT and *emx2^−/−^*. *emx2^−/−^* has 10-bp deletion compared to WT. Green denotes L and R TALEN target sites. Blue denotes BtsCI restriction site which is disrupted in *emx2^−/−^* embryos.

**Supplemental 2.**
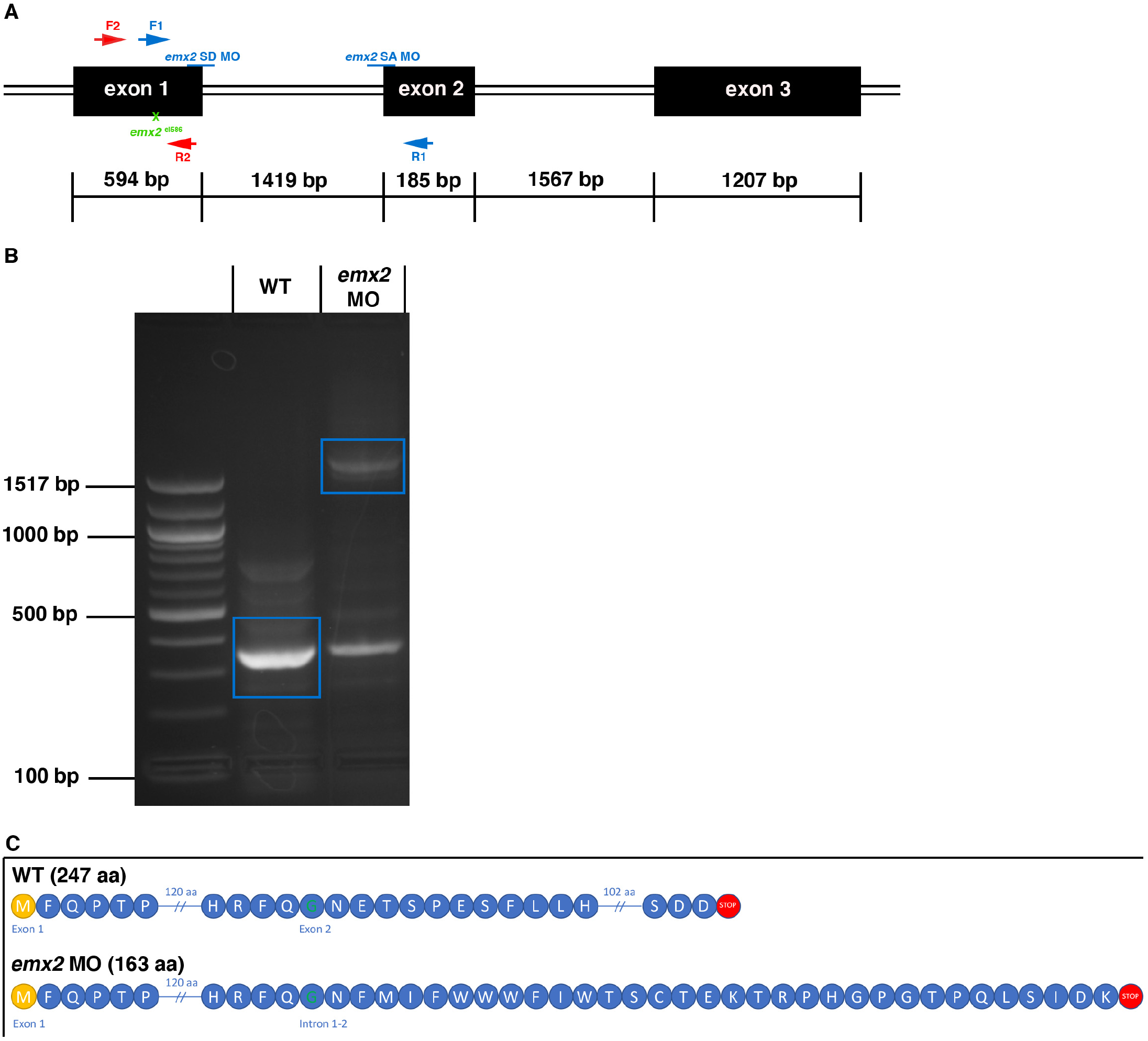
(A) Schematic of *emx2* DNA, where black boxes depict exons. Blue lines mark splice donor (SD) and splice acceptor (SA) MO. Green X marks site of *emx2^el586/el586^* TALEN disruption. Red and blue arrows mark forward and reverse primers binding sites. (B) Image of DNA product after RT-PCR. Control band is 350 bp in size, while *emx2* MO causes the appearance of a smaller band around 350 bp and a larger band at 1769 bp, indicating presence of intron 1. (C) Schematic of Emx2 protein after sequencing of WT and MO. The control Emx2 protein is 247 aa long. The Emx2 protein produced from *emx2* MO is truncated at 163 aa long as a result of inclusion of intron 1 between exon 1 and 2.

**Supplemental 3.**
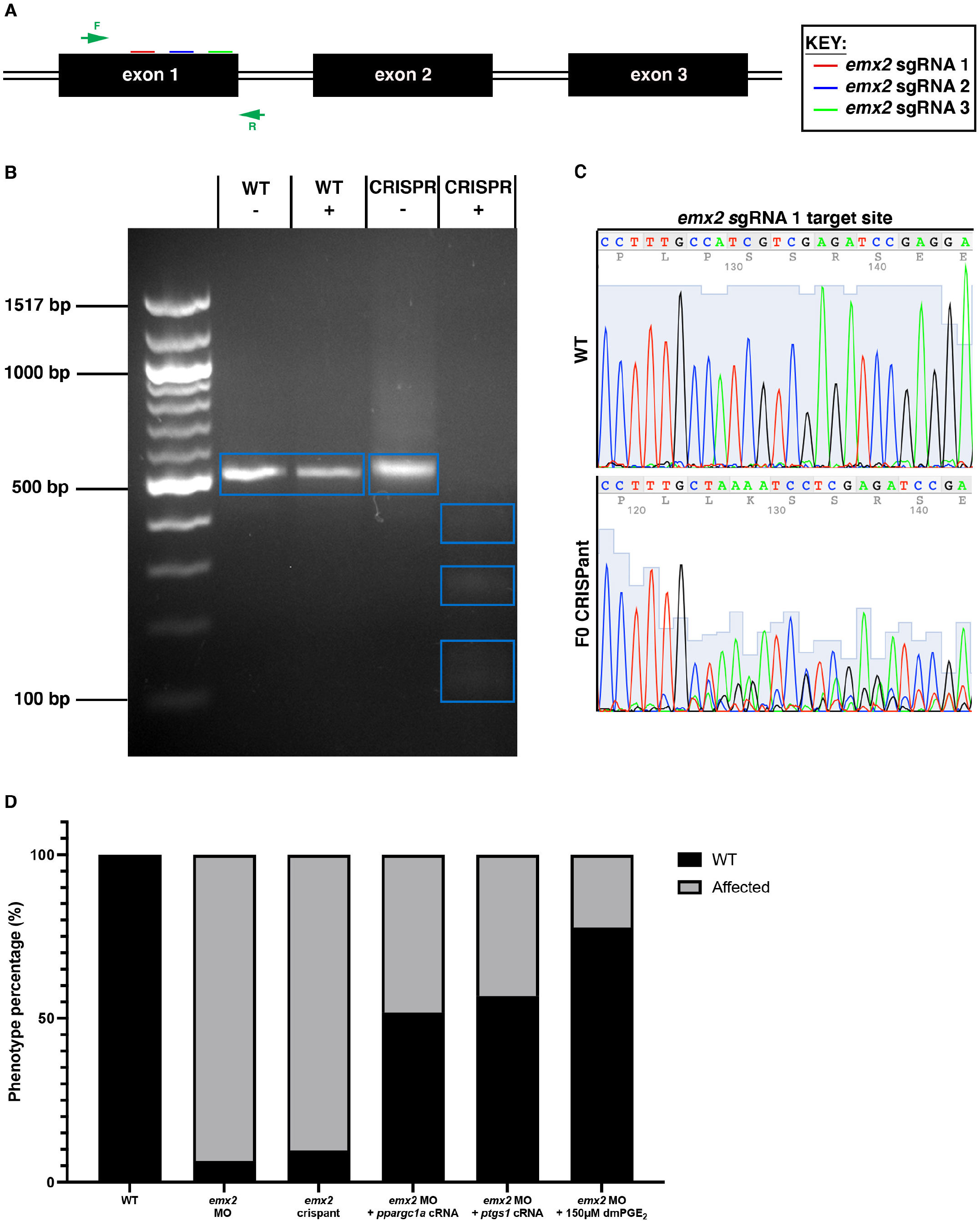
(A) Schematic of *emx2* DNA, where black boxes depict exons. Red, blue and green lines mark different *emx2* sgRNAs. Green arrows mark forward and reverse primers. (B) Image of DNA product after verifying crispants with T7 endonuclease assay. T7 endonuclease assay verifies cutting of *emx2* crispant samples. − denotes samples without T7 endonuclease, + denotes samples with T7 endonuclease. WT bands are present at 550 bp for both conditions. Crispant - shows similar presence to both WT conditions, while crispant + are cut into smaller bands with T7 endonuclease presence. (C) Sanger sequencing of *emx2* sgRNA AA target site between WT and F0 crispant. (D) Graph of phenotype percentage between WT and different experimental groups.

**Supplemental 4.**
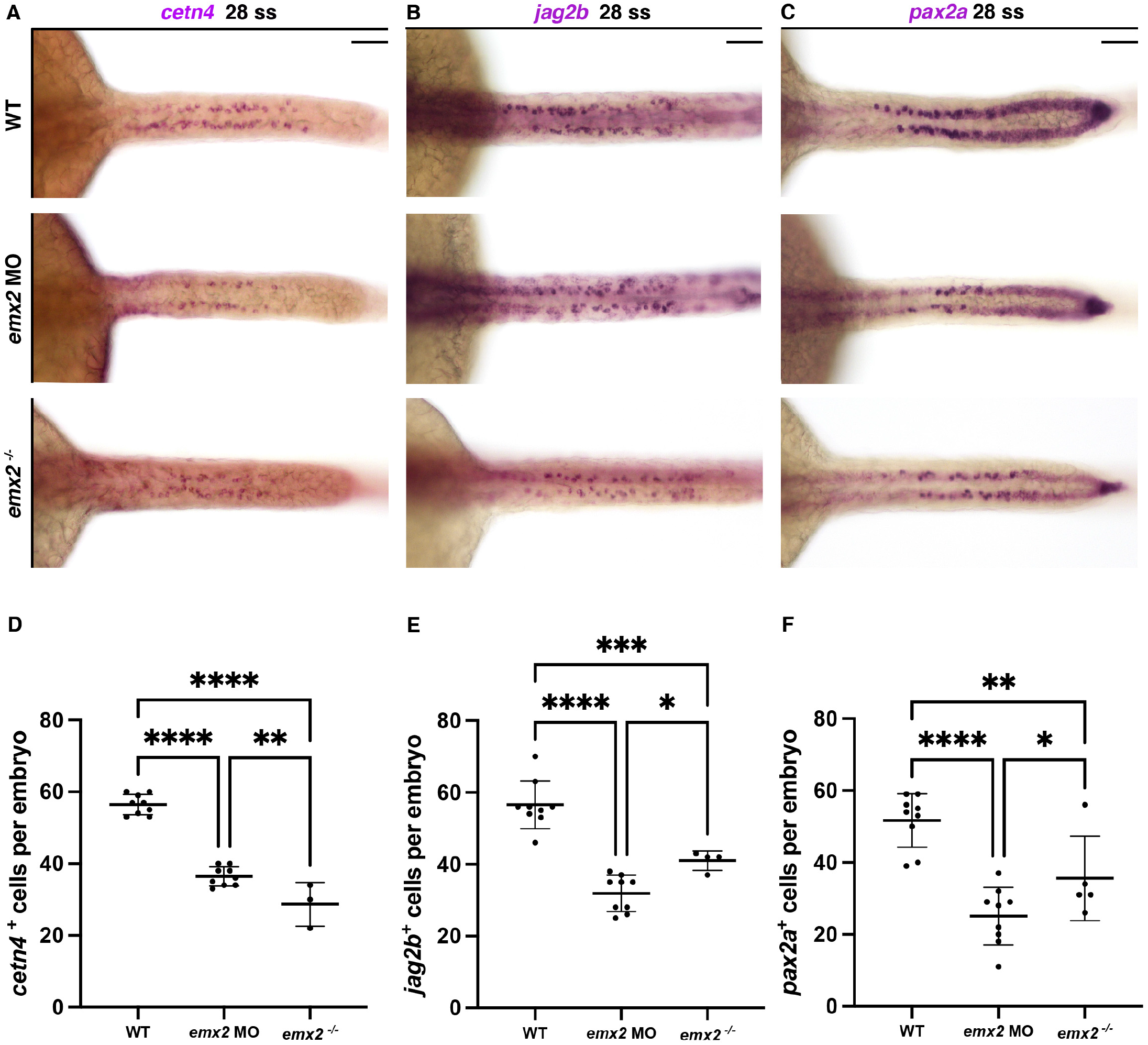
(A) 28 ss WT, *emx2* MO and *emx2^−/−^* embryos stained via WISH for *cetn4*. Scale bar = 50 μm. (B) 28 ss WT, *emx2* MO and *emx2^−/−^* embryos stained via WISH for *jag2b*. Scale bar = 50 μm. (C) 28 ss WT, *emx2* MO and *emx2^−/−^* embryos stained via WISH for *pax2a*. Scale bar = 50 μm. (D) *cetn4^+^* cells per embryo between 28 ss WT, *emx2* MO and *emx2^−/−^* embryos. (E) *jag2b^+^*cells per embryo between 28 ss WT, *emx2* MO and *emx2^−/−^* embryos. (F) *pax2a^+^* cells per embryo between 28 ss WT, *emx2* MO and *emx2^−/−^* embryos. Data presented on graphs are represented as mean ± SD; * p<0.05, ** p< 0.01 ***p < 0.001 and ****p < 0.0001 (t-test or ANOVA).

**Supplemental 5.**
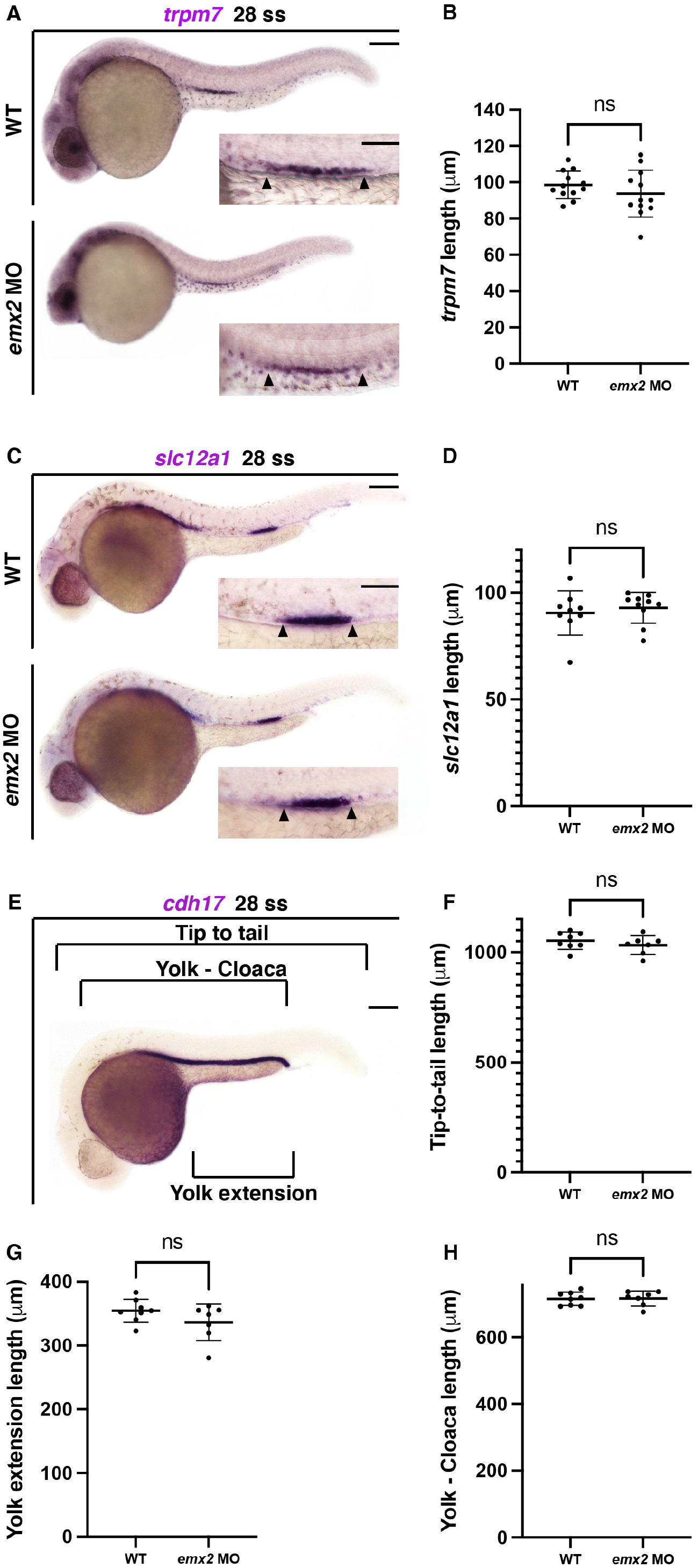
(A) 28 ss WT and *emx2* MO embryos stained via WISH for *trpm7.* Scale bars = 100 μm, inset = 50 μm. (B) *trpm7* length between 28 ss WT and *emx2* MO embryos. (C) 28 ss WT and *emx2* MO embryos stained via WISH for *slc12a1.* Scale bars = 100 μm, inset = 50 μm. (D) *slc12a1* length between 28 ss WT and *emx2* MO embryos. (E) Schematic of a 28 ss WT zebrafish embryo stained via WISH for *cdh17*. Black brackets denote the length of the measurement of each method. Scale bars = 100 μm. (F) Tip-to-tail length between 28 ss WT and *emx2* MO embryos. (G) Yolk extension length between 28 ss WT and *emx2* MO embryos. (H) Yolk-cloaca length between 28 ss WT and *emx2* MO embryos. Data presented on graphs are represented as mean ± SD; * p<0.05, ** p< 0.01 ***p < 0.001 and ****p < 0.0001 (t-test or ANOVA).

**Supplemental 6.**
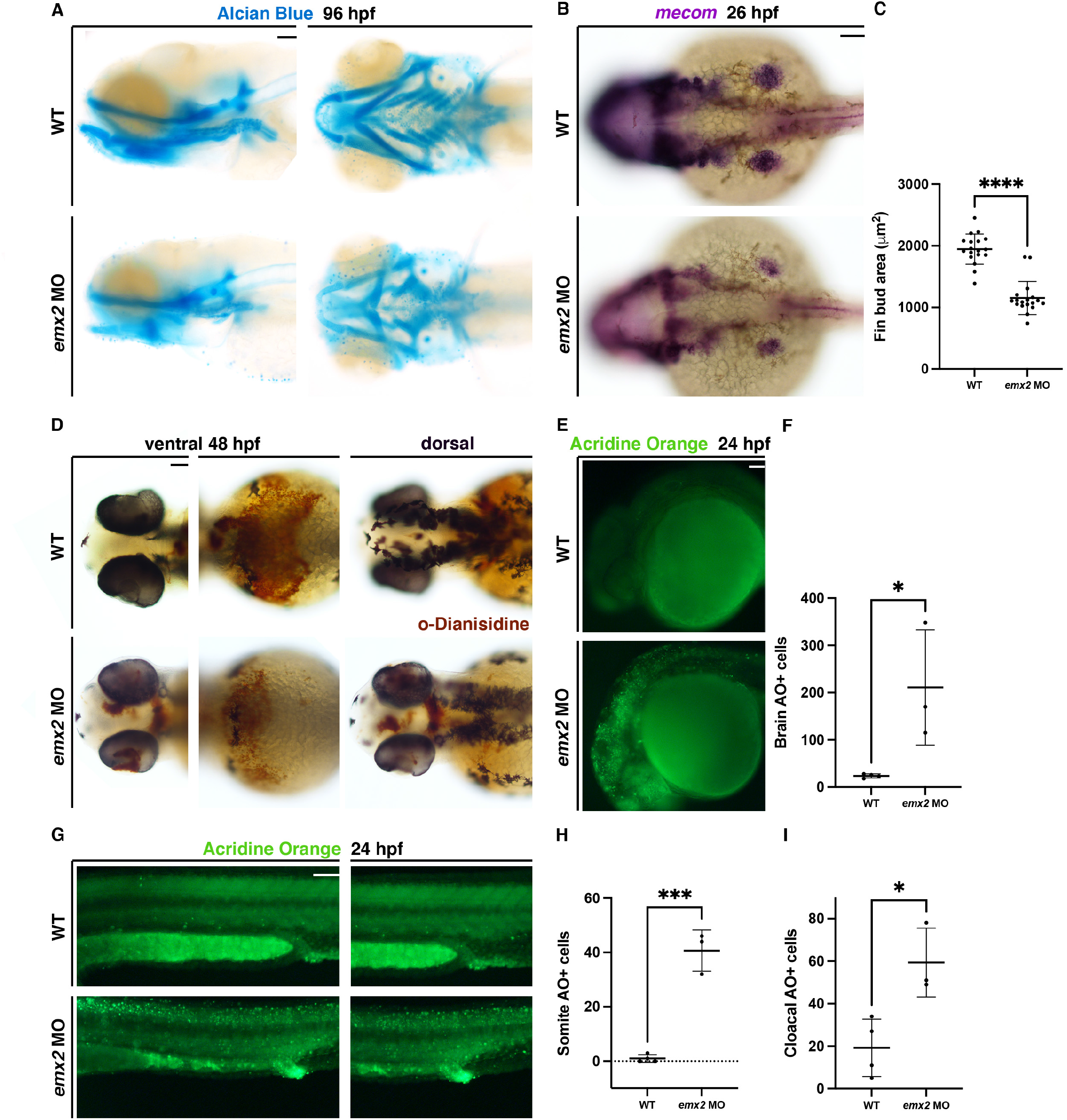
(A) 96 hpf Alcian Blue staining between WT and *emx2* MO embryos. Scale bar = 50 μm. (B) 26 hpf WT and *emx2* MO embryos stained via WISH for *mecom.* Scale bar = 50 μm. (C) Fin bud area between 26 hpf WT and *emx2* MO embryos. (D) 48 hpf WT and *emx2* MO embryos stained with o-Dianisidine to label erythrocytes. Scale bar = 50 μm. (E) 24 hpf Acridine Orange staining between WT and *emx2* MO embryos in the head. Scale bar = 50 μm. (F) Number of brain AO+ cells between 24 hpf WT and *emx2* MO embryos. (G) 24 hpf WT and *emx2* MO embryos stained with Acridine Orange along the trunk of embryos. Scale bar = 50 μm. (H) Number of somite AO+ cells between 24 hpf WT and *emx2* MO embryos. (I) Number of cloacal AO+ cells between 24 hpf WT and *emx2* MO embryos. Data presented on graphs are represented as mean ± SD; * p<0.05, ** p< 0.01 ***p < 0.001 and ****p < 0.0001 (t-test or ANOVA).

**Supplemental 7.**
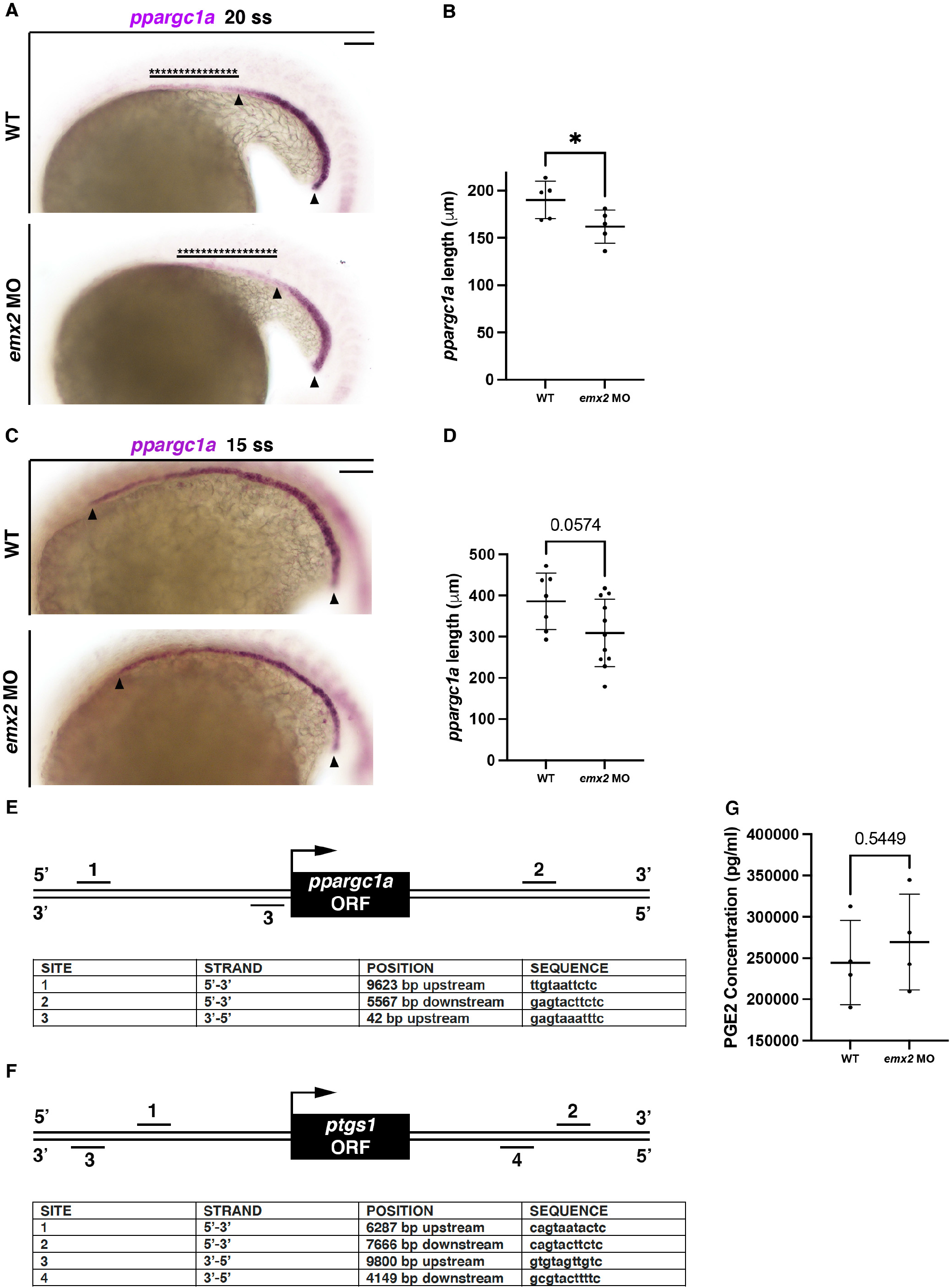
(A) 20 ss WT and *emx2* MO embryos stained via WISH for *ppargc1a.* Scale bar = 50 μm. (B) *ppargc1a* length between 20 ss WT and *emx2* MO embryos. (C) 15 ss WT and *emx2* MO embryos stained via WISH for *ppargc1a.* Scale bar = 50 μm. (D) *ppargc1a* length between 15 ss WT and *emx2* MO embryos. (E) Schematic of *ppargc1a* ORF with 10 kb upstream and downstream. Black lines denote conserved binding sites. (F) Schematic of *ptsg1* ORF with 10 kb upstream and downstream. Black lines denote conserved binding sites. (G) Relative concentration of PGE_2_ between 28 hpf WT and *emx2* MO embryos. Data presented on graphs are represented as mean ± SD; * p<0.05, ** p< 0.01 ***p < 0.001 and ****p < 0.0001 (t-test or ANOVA).

**Supplemental 8.**
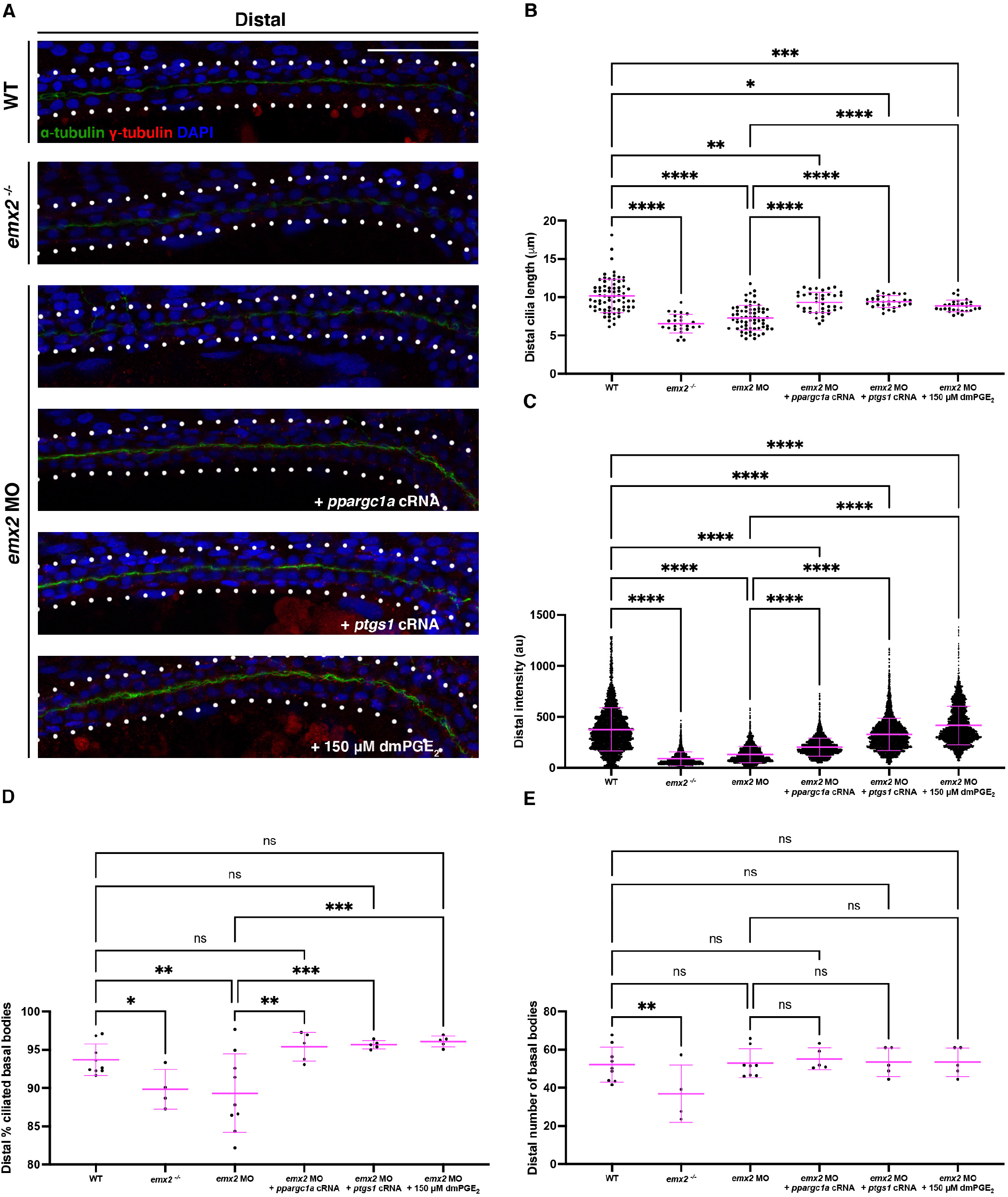
(A) 28 hpf whole-mount IF for acetylated α-tubulin (cilia, green), γ-tubulin (basal bodies, red), and DAPI (nucleus, blue) in the distal pronephros between different treatment groups. Scale bar = 50 μm. (B) Cilia length for the distal pronephros between different treatment groups. (C) Fluorescence intensity plot of α-tubulin intensity in the distal pronephros between different treatment groups. (D) Percentage of ciliated basal bodies/total basal bodies in distal pronephros between different treatment groups. (E) Number of basal bodies in the distal pronephros between different treatment groups. Data presented on graphs are represented as mean ± SD; * p<0.05, ** p< 0.01 ***p < 0.001 and ****p < 0.0001 (t-test or ANOVA).

